# Stability of working memory in continuous attractor networks under the control of short-term plasticity

**DOI:** 10.1101/424515

**Authors:** Alexander Seeholzer, Moritz Deger, Wulfram Gerstner

**Affiliations:** School of Computer and Communication Sciences and School of Life Sciences, Brain Mind Institute, École Polytechnique Fédérale de Lausanne, Lausanne, Switzerland; Institute for Zoology, Faculty of Mathematics and Natural Sciences, University of Cologne, Cologne, Germany

## Abstract

Continuous attractor models of working-memory store continuous-valued information in continuous state-spaces, but are sensitive to noise processes that degrade memory retention. Short-term synaptic plasticity of recurrent synapses has previously been shown to affect continuous attractor systems: short-term facilitation can stabilize memory retention, while short-term depression possibly increases continuous attractor volatility. However, it currently remains unclear to which degree these two short-term plasticity mechanisms interact, what their combined quantitative effect on working memory stability is, and whether these effects persist in neuronal networks with spike-based transmission. Here, we present a comprehensive description of the effects of short-term plasticity on noise-induced memory degradation in one-dimensional continuous attractor models. Our theoretical description, applicable to spiking and rate-based models alike, accurately describes the slow dynamics of stored memory positions in separate processes of diffusion due to spiking variability and drift due to sparse connectivity and neuronal heterogeneity. We find that facilitation decreases both diffusion and directed drifts, while short-term depression tends to increase both. Using mutual information, we evaluate the combined impact of short-term facilitation and depression on the ability of networks to retain stable working memory. Finally, our theory establishes links to experiments: we are able to predict the sensitivity of continuous working memory to distractor inputs and place constraints on network and synapse properties necessary to implement stable working memory.

**Author summary:** The ability to transiently memorize positions in the visual field is crucial for behavior. Models and experiments have shown that such memories can be maintained in networks of cortical neurons with a continuum of possible activity states, that reflects the continuum of positions in the environment. However, the accuracy of positions stored in such networks will degrade over time due to the noisiness of neuronal signaling and imperfections of the biological substrate. Previous work in simplified models has shown that synaptic short-term plasticity could stabilize this degradation by dynamically up- or down-regulating the strength of synaptic connections, thereby ”pinning down” memorized positions. Here, we present a general theory that accurately predicts the extent of this ”pinning down” by short-term plasticity in a broad class of biologically plausible models, thereby untangling the interplay of varying biological sources of noise with short-term plasticity. Importantly, our work provides a direct and novel theoretical link from the microscopic substrate of working memory – neurons and synaptic connections – to observable behavioral correlates. This allows us to constrain properties of cortical networks that are currently hard to assess experimentally, which we hope will help guide future theoretical and experimental work.

## Introduction

Information about past environmental stimuli can be stored and retrieved seconds later from working memory [1, 2]. Strikingly, this transient storage is achieved for timescales of seconds with neurons and synapse transmission operating mostly on time scales of tens of milliseconds and shorter [3]. An influential hypothesis of neuroscience is that working memory emerges from recurrently connected cortical neuronal networks: memories are retained by self-generating cortical activity through positive feedback [4–7], thereby bridging the time scales from milliseconds (neuronal dynamics) to seconds (behavior).

Sensory stimuli are often embedded in a physical continuum: for example, positions of objects in the visual field, frequencies of auditory stimuli, or the intensity and position of somatosensory stimuli on the body, all have continuously varying values. Ideally, the organization of cortical working memory circuits should reflect the continuous nature of sensory information [3]. A class of cortical working memory models able to store continuously structured information is that of *continuous attractors*, characterized by a continuum of meta-stable states, which can be used to retain memories over delay periods much longer than those of the single network constituents [8]. Continuous attractors were originally proposed as theoretical models for cortical working memory [9–11], path integration [12–14], and other cortical functions [15–17] (see e.g. [3, 18–21] for recent reviews), but recent experimental evidence for continuous attractor dynamics was indeed found in cortical networks [22], the limbic system [18, 23], and one-dimensional ring-attractors in the fly responsible for path integration and self-orientation [24, 25].

Continuous attractor models have been successfully employed in the context of visuospatial working memory to explain behavioral performance [26–29], to predict the effects of neuromodulation [30, 31], or the implications of cognitive impairment [32, 33]. However, when embedded in more realistic scenarios, the continuum of states quickly breaks down, since encoded memories are vulnerable to noise and heterogeneities that break, transiently or permanently, the crucial symmetry necessary for continuous attractors [10, 11, 13, 16, 34–40]. For example, the stochasticity of neuronal spiking (“fast noise”) leads to transient asymmetries that randomly displace encoded memories along the continuum of states [10, 11, 35, 37, 39, 40], leading, averaged over many trials, to *diffusion* of encoded information. More drastically, introducing fixed asymmetries (“frozen noise”) due to network heterogeneities causes a *directed drift* of memories and a collapse of the continuum of attractive states to a set of discrete states. Examples of heterogeneities in biological scenarios include the sparsity of recurrent connections [13, 36], or randomness in neuronal parameters [36] and values of recurrent weights [16, 34, 38]. Since both (fast) noise and heterogeneities are expected in cortical settings, the feasibility of continuous attractors as computational systems of the brain has been called into question [3, 6, 41].

Short-term plasticity of recurrent synaptic connections has been shown to influence the susceptibility of continuous attractor networks to both noise and heterogeneities. In particular, short-term depression has been observed to have strong effects on directed displacement of attractor states in rate models [42, 43], although no such effect was apparent in a spiking network implementation [44]. Short-term facilitation, on the other hand, has been proposed as a stabilizing mechanisms that could increase the retention time of memories in continuous attractor networks with noise-free rate neurons [38]. In simulations of a continuous attractor implemented with spiking neurons, facilitation with a fixed set of parameters was reported to cause slow drift [45, 46] and a reduced amount of diffusion [46]. However, despite the large number of existing studies, several fundamental questions remain unanswered. What are the quantitative effects of short-term facilitation in more complex neuronal models and across facilitation parameters? Does short-term depression influence the strength of diffusion and drift, and how does it interplay with facilitation? Do phenomena reported in rate networks persist in spiking networks? Finally, can a single theory be used to predict all of the effects observed in simulations?

Here, we present a comprehensive description of the effects of short-term facilitation and depression on noise-induced displacement of one-dimensional continuous attractor models. Extending earlier theories for diffusion [39, 40] and drift [38], we derive predictions of the amount of diffusion and drift in ring-attractor models with short-term plasticity. Our theory generalizes to a large class of neuron models, given their input-output relation, and to various biologically plausible sources of heterogeneity. The theoretical predictions are validated against a set of ring-attractor networks realized with spiking neurons for varying parameters of short-term facilitation and depression. We find that facilitation and depression play antagonistic roles: facilitation can decrease *both diffusion and drift* while depression *increases both*, which we show to have profound impact on the retention of memories, as measured by mutual information. Importantly, our theory is, to a large degree, independent of the microscopic network configurations, which allows relating it to experimentally observable quantities. We apply this insight to show that our theory can predict the sensitivity of networks with short-term plasticity to distractor stimuli. Finally, we demonstrate a second possible link to experiments by placing theoretical bounds on combinations of network and synapse properties that will yield stable working memory (as predicted by our model) under the presence of realistic biological variability.

## Results

We investigated, in theory and simulations, the effects of short-term synaptic plasticity (STP) on the dynamics of ring-attractor models consisting of *N* excitatory neurons with distance-dependent and symmetric excitation, and global (uniform) inhibition provided by a population of inhibitory neurons (Fig. 1A). For simplicity, we describe neurons in terms of firing rates, but our theory can be mapped to more complex neurons with spiking dynamics. An excitatory neuron *i* with 0 ≤ *i* < *N* is assigned an angular position 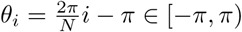, where we identify the bounds of the interval to form a ring topology (Fig. 1A). The firing rate *ϕ_i_* (in units of Hz) for each excitatory neuron *i* (0 ≤ *i* < *N* − 1) is given as a function of the neuronal input:

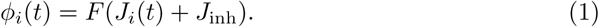

**Fig 1.**
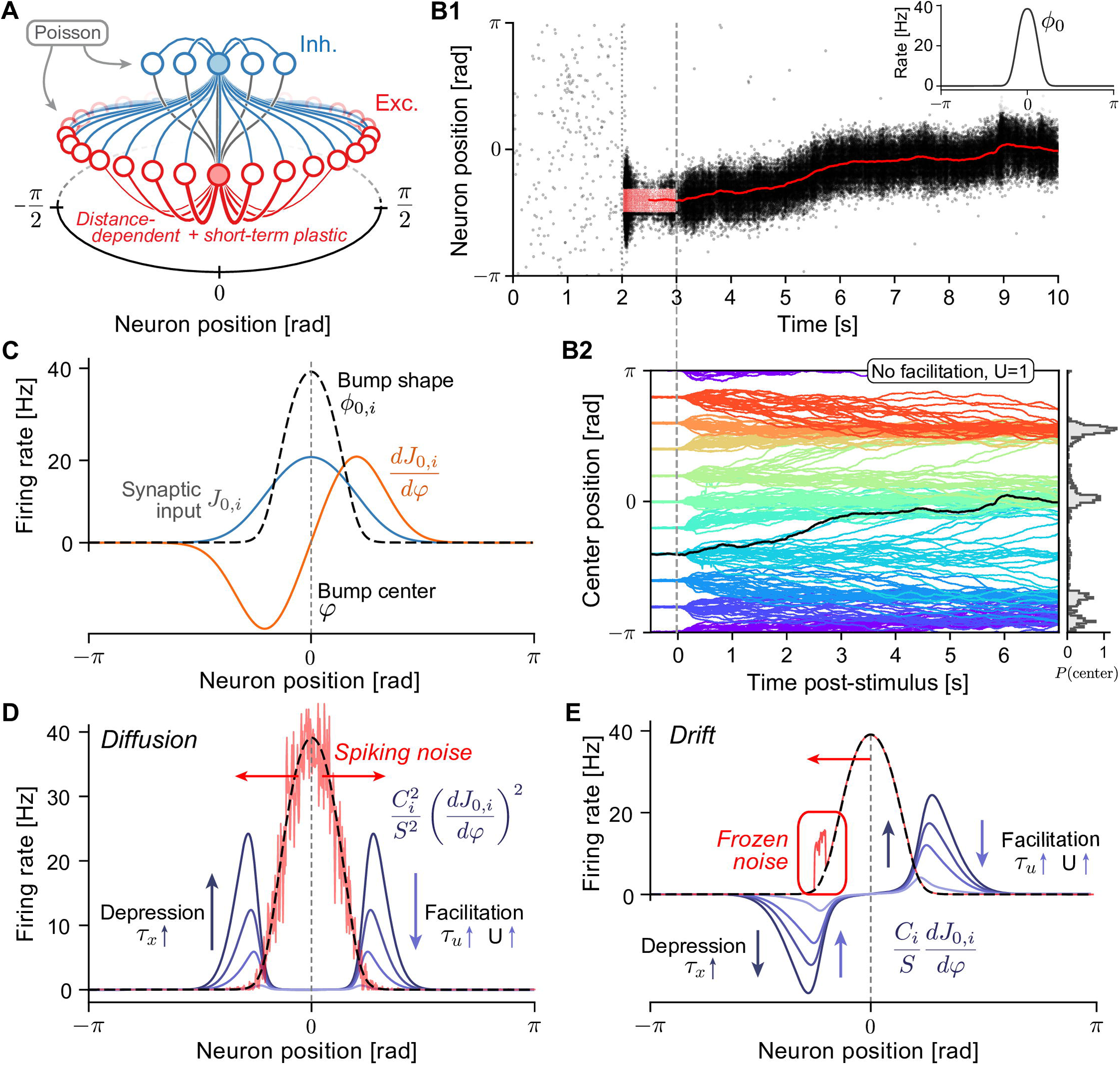
Drift and diffusion in ring-attractor models with short-term plasticity. **A** Excitatory (E) neurons (red circles) are distributed on a ring with coordinates in [−*π,π*]. Excitatory-to-excitatory (E-E) connections (red lines) are distance-dependent, symmetric, and subject to short-term plasticity (facilitation and depression, see Eq. (3)). Inhibitory (I) neurons (blue circles) project to all E and I neurons (blue lines) and receive connection from all E neurons (gray lines). Only outgoing connections from shaded neurons are displayed. In spiking simulations, neurons additionally receive excitatory input with spikes generated by homogeneous Poisson processes. **B1** Example simulation: E neurons fire asynchronously and irregularly at low rates until (dotted line) a subgroup of E neurons is stimulated (external cue), causing them to spike at elevated rates (red dots, input was centered at 0, starting at *t* = 2*s* for 1*s*). During and after (dashed line) the stimulus, a bump state of elevated activity forms and sustains itself after the external cue is turned off. The spatial center of the population activity is estimated from the momentary firing rates (red line, plotted from *t* = 2.5*s* onward). Inset: Activity profile in the bump state, centered at 0. **B2** Center positions of 20 repeated spiking simulations for 10 different initial cue positions each for a network with short-term depression (*U* = 1, *τ_x_* = 150*ms*). Random E-E connections (with connection probability *p* = 0.5) lead to directed drift in addition to diffusion. Right: Normalized histogram (200 bins) of final positions at time *t* = 13.5. **C** Illustration of quantities used in theoretic calculations. Neurons in the bump fire at rates *ϕ*_0,*i*_ (dashed black line, compare to B1, inset) due to the steady-state synaptic input *J*_0,*i*_ (blue line). Movement of the bump center causes a change of the synaptic input 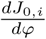 (orange line). **D** Diffusion along the attractor manifold is calculated (see Eq. (5)) as a weighted sum of the neuronal firing rates in the bump state (dashed black line). Spiking noise (red line) is illustrated as a random deviation from the mean rate with variance proportional to the rate. The symmetric weighting factors (blue lines show 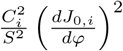 for varying *U*) are non-zero at the flanks of the firing rate profile. Stronger short-term depression and weaker facilitation increase the magnitude of weighting factors. **E** Deterministic drift is calculated as a weighted sum (see Eq. (7)) of systematic deviations of firing rates from the bump state (frozen noise): a large positive firing rate deviation in the left flank (red line) will cause movement of the center position to the left (red arrow) because the weighting factors (blue lines show 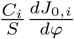 for varying *U*) are asymmetric.

Here, the input-output relation *F* relates the dimensionless excitatory *J_i_* and inhibitory *J*_inh_ inputs of neuron *i* to its firing rate. This represents a rate-based simplification of the possibly complex underlying neuronal dynamics [47]. We assume that the excitatory input *J_i_*(*t*) to neuron *i* at time *t* is given by a sum over all presynaptic neurons

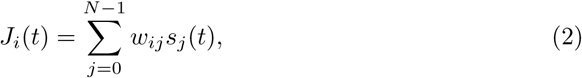

where *w_ij_s_j_*(*t*) describes the total activation of synaptic input from the presynaptic neuron *j* onto neurons *i*. The maximal strength *w_ij_* of recurrent excitatory-to-excitatory connections is chosen to be local in the angular arrangement of neurons, such that connections are strongest to nearby excitatory neurons (Fig. 1A, red lines). The momentary input depends also on the synaptic activation variables *s_j_*, to be defined below. Finally, connections to and from inhibitory neurons are assumed to be uniform and global (all-to-all) (Fig. 1A, blue lines), thereby providing non-selective inhibitory input *J*_inh_ to excitatory neurons.

As a model of STP, we assume that excitatory-to-excitatory connections are subject to short-term facilitation and depression, which we implemented using a widely adopted model of short-term synaptic plasticity [48]. The *outgoing synaptic activations s_j_* of neuron *j* are modeled by the following system of ordinary differential equations:

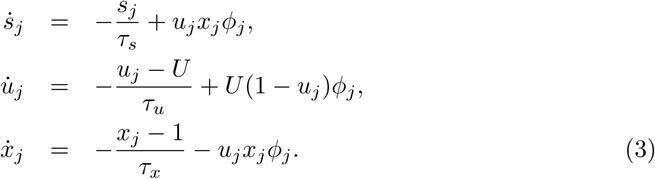

The timescale of recovery *τ_x_* is the main parameter of depression. While the recovery from facilitation is controlled by the timescale *τ_u_*, the parameter 0 < *U* ≤ 1 controls the baseline strength of unfacilitated synapses as well as the timescale of their strengthening. For fixed *τ_u_*, we consider smaller values of *U* to lead to a “stronger” effect of facilitation, and take *U* = 1 as the limit of non-facilitating synapses.

As a reference implementation of this model, we simulated networks of spiking conductance-based leaky-integrate-and-fire (LIF) neurons with (spike-based) short-term plastic synaptic transmission (Fig. 1B1, see *Spiking network model* in Materials and Methods for details). For these networks, under the assumption that neurons fire with Poisson statistics and the network is in a stationary state, neuronal firing can be approximated by the input-output relation *F* of Eq. (1) [49, 50] (see *Firing rate approximation* in Materials and Methods), which allows us to map the network into the general framework of Eqs. (1)-(2). In the stationary state, synaptic depression will lead to a saturation of the synaptic activation variables *s_j_* at a constant value as firing rates increase. This nonlinear behavior enables spiking networks to implement bi-stable attractor dynamics with relatively low firing rates [45, 51] similar to saturating NMDA synapses [11, 46]. Since we found that without depression (for *τ_x_* → 0) the bump state was not stable at low firing rates (in agreement with [51]), we always keep the depression timescale *τ_x_* at positive values.

Particular care was taken to ensure that networks display nearly identical bump shapes (similar to Fig. 1B1, inset; see also S1 Fig), which required the re-tuning of network parameters (recurrent conductance parameters and the width of distance-dependent connections; see *Optimization of network parameters* in Materials and Methods) for each combination of the STP parameters above.

Spiking simulations generally show a bi-stability between a *non-selective* state and a *bump* state. In the non-selective state, all excitatory neurons emit action potentials asynchronously and irregularly at roughly identical and low firing rates (Fig. 1B1, left of dotted line). The bump state can be evoked by stimulating excitatory neurons localized around a given position by additional external input (Fig. 1B1, red dots). After the external cue is turned off, a self-sustained firing rate profile (“bump”) emerges (Fig. 1B1, right of dashed line, and inset) that persists until the network state is again changed by external input. For example, a short and strong uniform excitatory input to all excitatory neurons causes a transient increase in inhibitory feedback that is strong enough to return the network to the uniform state [11].

During the bump state, fast fluctuations in the firing of single neurons transiently break the perfect symmetry of the firing rate profile and introduce small random displacements along the attractor manifold, which become apparent as a random walk of the center position. If the simulation is repeated for several trials, the bump has the same shape in each trial, but information on the center position is lost in a *diffusion*-like process. We additionally included varying levels of biologically plausible sources of heterogeneity (frozen noise) in our networks: *random connectivity* between excitatory neurons (E-E) and heterogeneity of the single neuron properties of the excitatory population [36], realized as a random distribution of leak reversal potentials. Heterogeneities makes the bump *drift* away from its initial position in a directed manner. For example, the bump position in the randomly connected (*p* = 0.5) network of Fig. 1B1 shows a clear upwards drift towards center positions around 0. Repeated simulations of the same attractor network with bumps initialized at different positions provide a more detailed picture of the combined drift and diffusion dynamics: bump center trajectories systematically are biased towards a few stable fixed points (Fig. 1B2) around which they are distributed for longer simulation times (histogram in Fig. 1B2, *t* = 13.5*s*). The theory developed in this paper aims at analyzing the above phenomena of drift and diffusion of the bump center.

### Theory of diffusion and drift with short-term plasticity

To untangle the observed interplay between diffusion and drift and investigate the effects of short-term plasticity, we derived a theory that reduces the microscopic network dynamics to a simple one-dimensional stochastic differential equation for the bump state. The theory yields analytical expressions for diffusion coefficients and drift fields, that depend on short-term plasticity parameters, the shape of the firing rate profile of the bump, as well as the neuron model chosen to implement the attractor.

First, we *assume* that the system of Eqs. (3) together with the network Eqs. (1)-(2) has a 1-dimensional manifold of meta-stable states, i.e. the network is a ring-attractor network as described in the introduction. This entails, that the network dynamics permit the existence of a family of solutions that can be described as a self-sustained and symmetric bump of firing rates *ϕ*_0,*i*_(*φ*) = *F*(*J*_0,*i*_(*φ*)) with corresponding inputs *J*_0,*i*_(*φ*) (for 0 ≤ *i* < *N*). Importantly, the center *φ* of the bump can be located at any arbitrary position 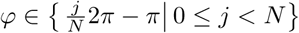. For example, if *ϕ*_0,*i*_(0) is a solution with input *J*_0,*i*_(0), then 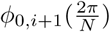 is also a solution with input 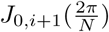. This solution is illustrated in Fig. 1C for a bump centered at *φ* = 0. Second, we assume that the number *N* of excitatory neurons is large (*N* → ∞), such that we can think of the possible positions *φ* as a continuum. Our final assumption is that neuronal firing is noisy, with spike counts distributed as Poisson processes, and that we are able to replace the shot-noise of Poisson spiking by white Gaussian noise with the same mean and autocorrelation (see *Diffusion* in Materials and Methods, and Discussion). Under these assumptions, we are able to reduce the network dynamics to a **one-dimensional Langevin equation**, describing the dynamics of the center *φ*(*t*) of the firing rate profile (see *Analysis of drift & diffusion with STP* in Materials and Methods):

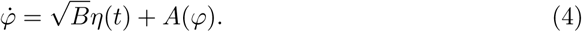

Here, *η*(*t*) is white Gaussian noise with zero mean and correlation function 〈*η*(*t*), *η*(*t*′)〉 = · (*t* – *t*′).

The first term is diffusion characterized by a ***diffusion strength** B*^1^, which describes the random displacement of bump center positions due to fluctuations in neuronal firing. For *A*(*φ*) = 0 this term causes diffusive displacement of the center *φ*(*t*) from its initial position *φ*(*t*_0_), with a mean (over realizations) squared displacement of positions 〈[*φ*(*t*) − *φ*(*t*_0_)]^2^〉 = *B* · (*t* – *t*_0_) that, during an initial phase, increases linearly with time [14, 52, 53], before saturating due to the circular domain of possible center positions [39]. Our theory shows (see *Diffusion* in Materials and Methods) that the coefficient *B* can be calculated as a weighted sum over the neuronal firing rates (Fig. 1D)

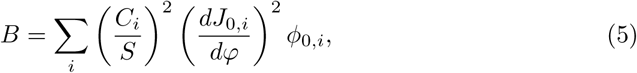

where 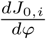 is the change of the input to neuron *i* under shifts of the center position (Fig. 1C, orange line), and *S* is a normalizing constant.

The analytical factors *C_i_* express the spatial dependence of the diffusion coefficient on the short-term plasticity parameters through

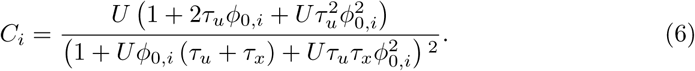

The dependence of the single summands in Eq. (5) on short-term plasticity parameters is visualized in Fig. 1D, where we see that: a) due to the squared spatial derivative 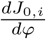 of the bump shape and the squared factors *C_i_*/*S*, the important contributions to the sum arise primarily from the flanks of the bump; b) for a fixed bump shape, summands increase with stronger short-term depression (larger *τ_x_*) and decrease with stronger short-term facilitation (smaller *U*, larger *τ_u_*). This result extends the approach of [39] to synaptic dynamics with short-term plasticity: the limiting case of no facilitation and depression (*U* = 1, *τ_x_* = 0*ms*) simplifies the normalization factor *S* considerably, leaves *C_i_* = 1, and recovers the result for diffusion the stated there [39, Eq. S18].

The second term in Eq. (5) is the ***drift field*** *A*(*φ*), which describes deterministic drifts due to the inclusion of heterogeneities. For heterogeneity caused by variations in neuronal reversal potentials and random network connectivity, we calculate (see *Frozen noise* in Materials and Methods) systematic deviations Δ*ϕ_i_*(*φ*) of the single neuronal firing rates from the steady-state bump shape that depend on the current position *φ* of the bump center. In *Drift* in Materials and Methods, we show that the drift field is then given by a weighted sum over the firing rate deviations:

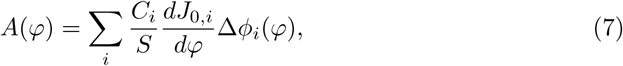

with weighing factors depending on the spatial derivative of the bump shape 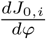 and the parameters of the synaptic dynamics through the same factors *C_i_*/*S*. This is illustrated in Fig. 1E: in contrast to Eq. (5) summands are now asymmetric with respect to the bump center, since the spatial derivative is not squared.

### Prediction of continuous attractor dynamics with short-term plasticity

To demonstrate the accuracy of our theory, we chose random connectivity as a first source of frozen variability. Random connectivity was realized in simulations by retaining only a random fraction 0 < *p* ≤ 1 (connection probability) of excitatory-to-excitatory (EE) connections. The uniform connections from and to inhibitory neurons are taken as all-to-all, since the effects of making these random and sparse would have only indirect effects on the dynamics of the bump center positions.

Our theory accurately predicts the drift-fields *A*(*φ*) (see Eq. (7)) induced by frozen variability in networks with short-term plasticity (Fig. 2). Briefly, for each neuron 0 ≤ *i* < *N*, we treat each realization of frozen variability as a perturbation Δ_*i*_ around the perfectly symmetric system and use an expansion to first order of the input-output relation *F* to calculate the resulting changes in firing rates (see *Frozen noise* for details):

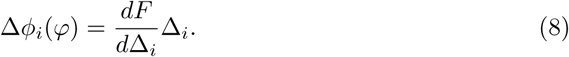

**Fig 2.**
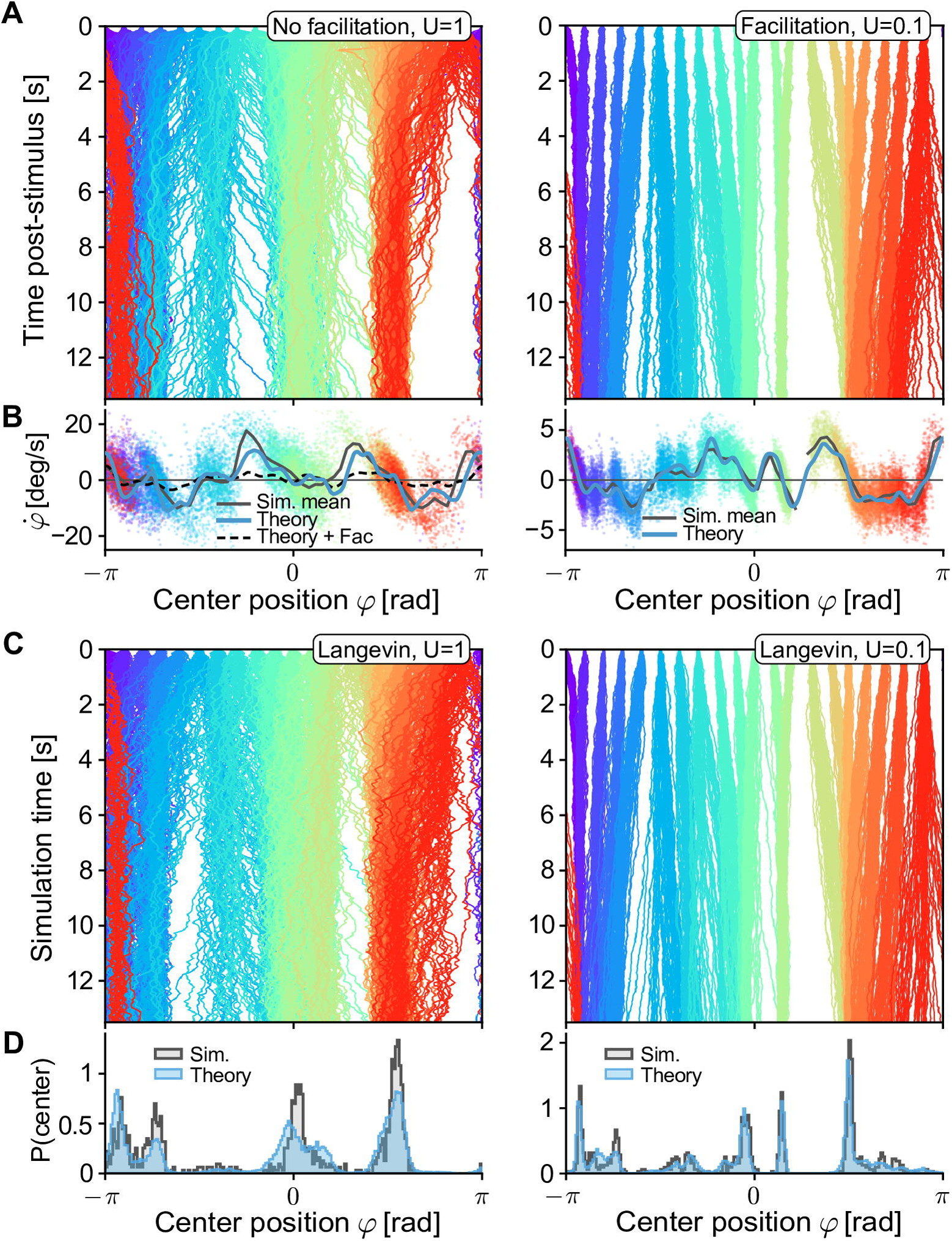
Drift field predictions for varying short-term facilitation. All networks have the same instantiation of random connectivity (*p* = 0.5), similar to Fig. 1B1. **A** Centers of excitatory population activity for 50 repetitions of 13.5*s* delay activity, for 20 different positions of initial cues (cue is turned off at *t* = 0) colored by position of the cues. Left: no facilitation (*U* = 1). Right: with facilitation (*U* = 0.1). **B** Drift field as a function of the bump position. The theoretical prediction (blue line, see Eq. (7)) of the drift field is compared to velocity estimations along the trajectories shown in A, colored by the line they were estimated from. The thick black line shows the binned mean of data points in 60 bins. For comparison, the predicted drift field for *U* = 0.1 is plotted (thin dashed line). Left: no facilitation (*U* = 1), for comparison the theoretical prediction for the case *U* = 0. 01 is plotted as a dashed line. Right: with facilitation (*U* = 0.01). **C** Trajectories under the same conditions as in A, but obtained by forward-integrating the one-dimensional Langevin equation, Eq. (4). **D** Normalized histograms of final positions at time *t* = 13.5 for data from spiking simulations (gray areas, data from A) and forward solutions of the Langevin equations (blue areas, data from C). Other STP parameters were: *τ_u_* = 650*ms, τ_x_* = 150*ms*.

The resulting terms are then used in Eq. (7) to predict the magnitude of the drift field *A*(*φ*) for any center position *φ*, which will, importantly, depend on STP parameters. The same approach can be used to predict drift fields induced by heterogeneous single neuron parameters [36] (see next sections) and additive noise on the E-E connection weights [16, 38].

We first simulated spiking networks with only short-term depression and without facilitation (Fig. 2A, left, same network as in Fig. 1B1), for one instantiation of random random (*p* = 0.5) connectivity. Numerical estimates of the drift in spiking simulations (by measuring the displacement of bumps over time as a function of their position, see *Spiking simulations* in Materials and Methods for details) yielded drift-fields in good agreement with the theoretical prediction (Fig. 2B, left). At points where the drift field prediction crosses from positive to negative values (e.g. Fig. 2B, left, 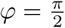), we expect stable fixed points of the center position dynamics in agreement with simulation results, which show trajectories converging to these points. Similarly, unstable fixed points (negative-to-positive crossings) can be seen to lead to a separation of trajectories (e.g. Fig. 2A, left, 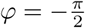). In regions where the positional drifts are predicted to lie close to zero (e.g. Fig. 2A, left *φ* = 0) the effects of diffusive dynamics are more pronounced. Finally, numerical integration of the full 1-dimensional Langevin equation Eq. (4) with coefficients predicted by Eqs. (5)-(7), produces trajectories with dynamics very similar to the full spiking network (Fig. 2C, left). When comparing the center positions after 13.5*s* of delay activity between the full spiking simulation and the simple 1-dimensional Langevin system, we found very similar distributions of final positions (Fig. 2D, left, compare to Fig. 1B1, histogram). Our theory thus produces an accurate approximation of the dynamics of center positions in networks of spiking neurons with STP, thereby reducing the complex dynamics of the whole network to a simple equation. It should be noted that, in regions with strong drift or steep negative-to-positive crossings, the numerically estimated drift-fields deviate from the theory due to under-sampling of these regions as trajectories move quickly through them, yielding fewer data points. In *Short-term plasticity controls drift* we show that for stronger heterogeneities the theory tends to generally over-predict drift-fields.

Introducing strong short-term facilitation (*U* = 0.1) reduces the predicted drift fields (Fig. 2B, left, dashed line), which resemble a scaled-down version of the drift-field for the unfacilitated case. We confirmed this theoretical prediction by simulations including facilitation (Fig. 2A, right): the resulting drift fields show significant reduction of speeds (Fig. 2B, right) while zero crossings remained similar to the unfacilitated network, similar to the results in [38]. Theoretical predictions of the drift fields with bump shapes extracted from these simulations again show an accurate prediction of the dynamics (Fig. 2B, right). Thus, as before, forward integrating the simple 1-dimensional Langevin-dynamics yields trajectories (Fig. 2C, right) highly similar to those of the full spiking network, with closely matching distributions of final positions (Fig. 2D, right), indicative of a matching strength of diffusion. In summary, our theory predicts the effects of STP on the joint dynamics of diffusion and drift due to network heterogeneities, which we will show in detail in the next sections.

### Short-term plasticity controls diffusion

To isolate the effects of STP on diffusion, we simulated networks *without frozen noise* for various STP parameters. For each combination of parameters, we simulated 1000 repetitions of 13.5s delay activity (after cue offset) distributed across 20 uniformly spaced initial cue positions (see Fig. 3A for an example). From these simulations, the strength of diffusion was estimated by measuring the growth of variance (over repetitions) of the distance of the center position from its initial position as a function of time (see *Spiking simulations* in Materials and Methods for details). For all parameters considered, this growth was well fit by a linear function (e.g. Fig. 3A, inset), the slope of which we compared to the theoretical prediction obtained from the diffusion strength *B* (Eq. (5)).

**Fig 3.**
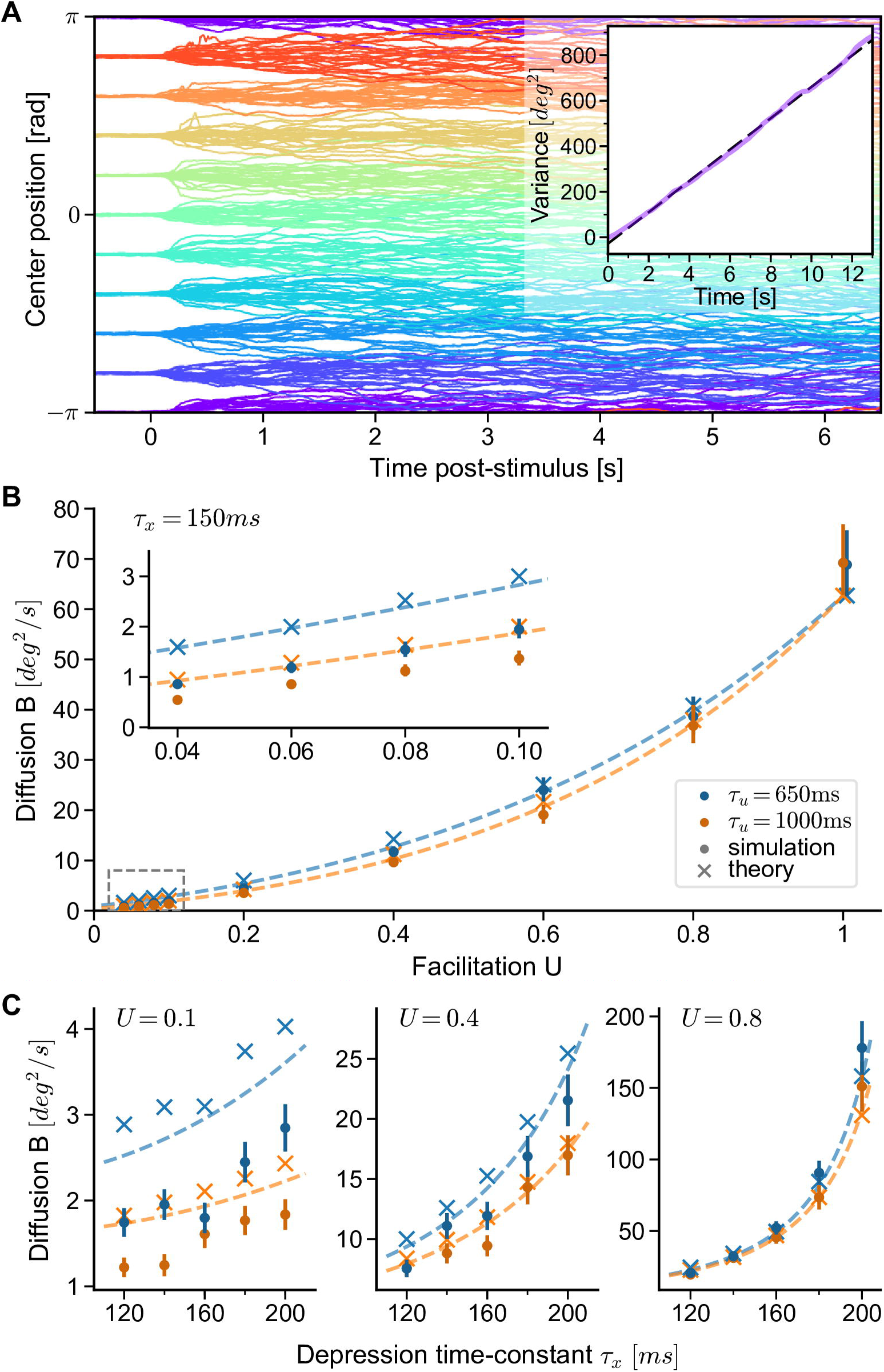
Diffusion on continuous attractors is controlled by short-term plasticity. **A** Center positions of 20 repeated simulations of the reference network (*U* = 1, *τ_x_* = 150ms) for 10 different initial cue positions each. Inset: Estimated variance of deviations of center positions *φ*(*t*) from their positions *φ*(0.5) at *t* = 0.5*s* (purple) as a function of time (〈[*φ*(*t*) – *φ*(0.5)]^2^〉), together with linear fit (dashed line). The slope of the dashed line yields an estimate of *B* (Eq. (5)). **B,C** Diffusion strengths estimated from simulations (dots, error bars show 95% confidence interval, estimated by bootstrapping) compared to theory. Dashed lines show theoretical prediction using firing rates measured from the reference network (*U* = 1, *τ_x_* = 150*ms*), while crosses are theoretical estimates using firing rates measured for each set of STP parameters separately (crosses). **B** Diffusion strength as a function of facilitation parameter *U*. Inset shows zoom of region indicated in the dashed area in the lower left. Increasing the facilitation time constant *τ_u_* = 650*ms* (blue) to *τ_u_* = 1*s* (orange) affects diffusion only slightly. In panels A and B, the depression time constant is *τ_x_* = 150*ms*. **C** Diffusion strength as a function of depression time constant *τ_x_*. Results for three different values of *U* are shown (note the change in scale). Colors indicate the two different values for the facilitation time constant also used in panel B.

We find that facilitation and depression control the amount of diffusion along the attractor manifold in an antagonistic fashion (Fig. 3B,C). First, increasing facilitation by lowering the facilitation parameter *U* from its baseline *U* = 1 (no facilitation) towards *U* = 0, while keeping the depression time constant *τ_x_* = 150*ms* fixed, decreases the measured diffusion strength over an order of magnitude (Fig. 3B, dots). On the other hand, increasing the facilitation time constant *τ_u_* from *τ_u_* = 650*ms* to *τ_u_* = 1000*ms* (Fig. 3B, orange and blue dots, respectively) only slightly reduces diffusion. Our theory further predicts that increasing the facilitation time constants above *τ_u_* = 1*s* will not lead to large reductions in the magnitude of diffusion (see S2 Fig). Second, we find that increasing the depression time constant *τ_x_* for fixed *U*, thereby slowing down recovery from depression, leads to an increase of the measured diffusion (Fig. 3C). More precisely, increasing the depression time constant from *τ_x_* = 120*ms* to *τ_x_* = 200*ms* leads only to slight increases in diffusion for strong facilitation (*U* = 0.1), but to a much larger increase for weak facilitation (*U* = 0.8).

For a comparison of these simulations with our theory, we used two different approaches. First, we estimated the diffusion strength by using the precise shape of the stable firing rate profile extracted separately for each network with different sets of parameters. This first comparison with simulations confirms that the theory closely describes the dependence of diffusion on short-term plasticity for each parameter set (Fig. 3B, crosses). The observed effects could arise directly from changes in STP parameters for a fixed bump shape, or indirectly since STP parameters also influence the shape of the bump. To separate such direct and indirect effects, we used for a second comparison a theory with fixed bump shape, i.e. the bump shape measured in a “reference network” (*U* = 1, *τ_x_* = 150*ms*) and extrapolated curves by changing only STP parameters in Eq. (5). This leads to very similar predictions (Fig. 3B, dashed lines) and supports the following conclusions: a) the diffusion to be expected in attractor networks with similar observable quantities (mainly, the bump shape) depends only on the short-term plasticity parameters; b) the bump shapes in the family of networks we have investigated are sufficiently similar to be approximated by measurement in a single reference network. It should be noted that the theory tends to slightly over-estimate the amount of diffusion, especially for small facilitation *U* (see Fig. 3B, C left). This may be because slower bump movement decreases the firing irregularity of flank neurons, which deviates from the Poisson firing assumption of our theory (see also Discussion). However, given the simplifying assumptions needed to derive the theory, the match to the spiking network is surprisingly accurate.

### Short-term plasticity controls drift

Having established that our theory is able to predict the effect of STP on diffusion, as well as drift for a single instantiation of random connectivity, we wondered how different sources of heterogeneity (frozen noise) would influence the drift of the bump. We considered two sources of heterogeneity: First, random connectivity as introduced above, and second, heterogeneity of the leak reversal potential parameters of excitatory neurons: leak reversal potentials of excitatory neurons are given by *V_L_* + Δ*_L_*, where Δ*_L_* is normally distributed with zero mean and standard deviation *σ_L_* [36]. The resulting fields can be calculated by calculating the resulting perturbations to the firing rates of neurons by Eq. (8) (see *Frozen noise* in Materials and Methods for details).

The theory developed so far allowed us to predict drift-fields for a given realization of frozen noise, controlled by the noise parameters *p* (for random connectivity) and *σ_L_* (for heterogeneous leak reversal-potentials) (see S3 Fig for a comparison of predicted drift fields to those measured in simulations for varying STP parameters and varying strengths of frozen noises). We wondered, whether we could take the level of abstraction of our theory one step further, by predicting the magnitude of drift fields from the frozen noise parameters only, independently of a specific realization. First, the expectation of drift fields under the distributions of the frozen noises vanishes for any given position: 〈*A* (*φ*)〉_frozen_ = 0, where the expectation 〈·〉_frozen_ is taken over both noise parameters. We thus turned to the expected squared magnitude of drift fields under the distributions of these parameters (see *Squared field magnitude* in Materials and Methods for the derivation):

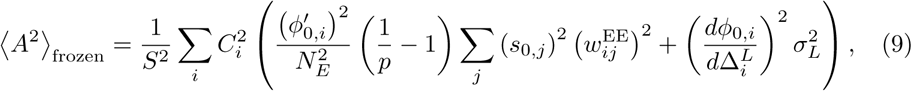

where *s*_0,*j*_ is the steady-state synaptic activation. Here, we introduced the derivatives of the input-output relation with respect to the noise sources that appear in Eq. (8): 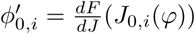 is the derivative with respect to the steady state synaptic input, and 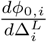 is the derivative with respect to the perturbation in the leak potential. In *Squared field magnitude* in Materials and Methods, we show that Eq. (9) is independent of the center position *φ*, and can be estimated from simulations as the variance of the drift field across positions, averaged over an ensemble of network instantiations.

We defined the root of the expected squared magnitude of Eq. (9) as the *expected field magnitude*:

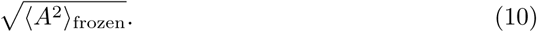

This quantity predicts the magnitude of the deviations of drift-fields from zero that are expected from the parameters that control the frozen noise – in analogy to the standard deviation for random variables, it predicts the standard deviation of the fields. To compare this quantity to simulations, we varied both heterogeneity parameters. First, the connectivity parameter *p* was varied between 0.25 and 1. Second, for heterogeneities in leak reversal-potentials, we chose values for the standard deviation *σ_L_* of leak-reversal potentials between 0*mV* and 1.5*mV*, which lead to a similar range of drift magnitudes as those of randomly connected networks. For each combination of heterogeneities and STP parameters (networks had either random connections or heterogeneous leaks) we then realized 18 – 20 networks, for which we simulated 400 repetitions of 6.5s of delay activity each (20 uniformly spaced positions of the initial cue). We then estimated the drift-field numerically by recording displacements of bump centers along their trajectories (as in Fig. 2A, B) and measured the standard deviation of the resulting fields across all positions.

Similar to the analysis of diffusion above, we find that facilitation and depression elicit antagonistic control over the magnitude of drift fields. In both simulations and theory, we find (Fig. 4A,B) that the expected field magnitude *decreases* as the effect of facilitation is *increased* from unfacilitated networks (*U* = 1) through intermediate levels of facilitation (*U* = 0.4) to strongly facilitating networks (*U* = 0.1). Our theory predicts this effect surprisingly well, which we validated twofold (as for the diffusion magnitude). First, we used Eq. (10) with all parameters and coefficients estimated from each spiking simulation separately (Fig. 4A,B, plus-signs and crosses). Second, we extrapolated the theoretical prediction by using coefficients in Eq. (9) measured from the unfacilitated reference network only (*U* = 1, *τ_x_* = 150*ms*) but changed the facilitation and heterogeneity parameters (Fig. 4A,B, dashed lines). The largest differences between the extrapolated and full theory are seen for *U* < 1 and randomly connected networks (*p* < 1), which we found to result from the fact that bump shapes for these networks tended to be slightly reduced under random and sparse connectivity (e.g. the top firing rate is reduced to ~35*Hz* for *U* = 0.1, *p* = 0.25). Generally, as noise levels increase, our theory tends to over-estimate the squared magnitude of fields, since we rely on a linear expansion of perturbations to the firing rates to calculate fields (Eq. (8)). Such deviations are expected as the magnitude of firing rate perturbations increases, and could be counter-acted by including higher-order terms. Since in the theory facilitation (and depression) only scales the firing rate perturbations (Eq. (7)), these deviations can also be observed across facilitation parameters. Finally, we performed a similar analysis to investigate the effect of short-term depression on drift fields. Here, we varied the depression time constant *τ_x_* for randomly connected networks with *p* = 0.6, by simulating networks with combinations of short-term plasticity parameters from *U* ∈ {0.1,0.4, 0.8} and *τ_x_* ∈ {120*ms*, 160*ms*, 200*ms*} (Fig. 4C). We find that an increase of the depression time constant leads to increased magnitude of drift fields, which again is well predicted by our theory.

**Fig 4.**
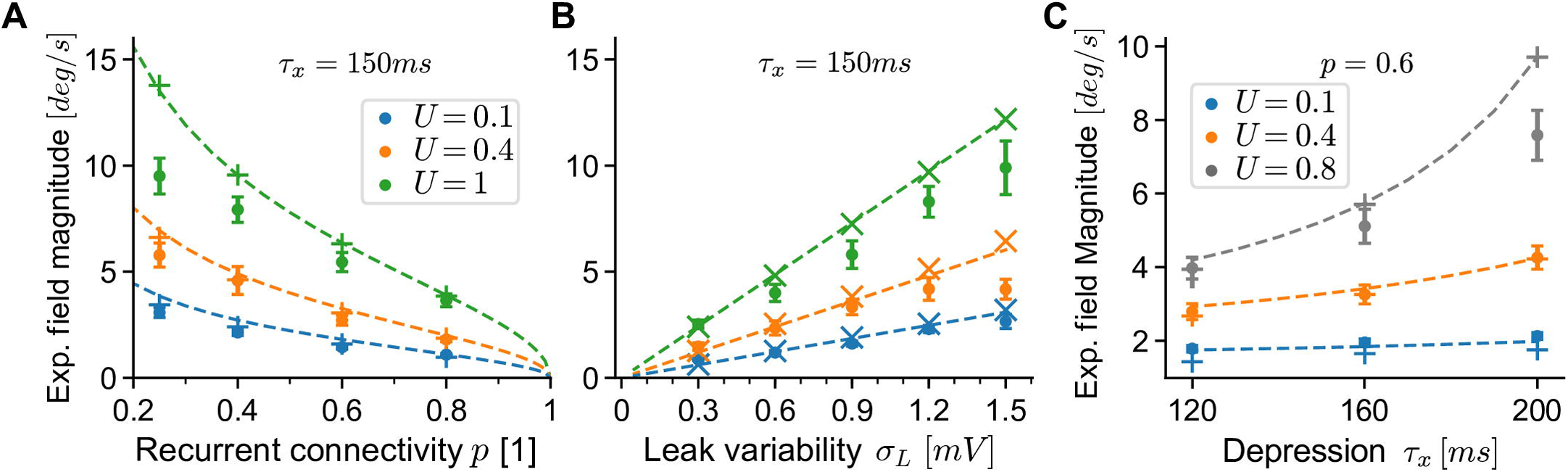
Drift field magnitude is controlled by short-term plasticity. **A** Expected magnitude of drift fields as a function of the sparsity parameter *p* of recurrent excitatory-to-excitatory connections. Dots are the standard deviation of fields estimated from 400 trajectories (see main text) of each network, averaged over 18 – 20 realizations for each noise parameter and facilitation setting (error bars show 95% confidence of the mean). Theoretical predictions (dashed lines) are given by Eq. (10) extrapolated from the reference network (*U* = 1, *τ_x_* = 150). For validation, we also estimated Eq. (10) with coefficients measured from each simulated network separately (plus signs). The depression time constant was *τ_x_* = 150ms. **B** Same as in panel A, with heterogeneous leak-reversal potentials as the source of frozen noise. Validation predictions are plotted as crosses. **C** Same as in panels A,B but varying the depression time constant *τ_x_* for a fixed level of frozen noise (random connectivity, *p* = 0.6). In all panels, the facilitation time constant was *τ_u_* = 650ms.

### Short-term plasticity controls memory retention

The theory developed in previous sections shows that diffusion and drift of the bump center *φ* are controlled antagonistically by short-term depression and facilitation. In a working memory setup, we can view the attractor dynamics as a noisy communication channel [54] that maps a set of initial positions *φ*(*t* = 0*s*) (time of the cue offset in the attractor network) to associated final positions *φ*(*t* = 6.5*s*), after a memory retention delay of 6.5*s*. We used the distributions of initial and (associated) final positions to investigate the combined impact of diffusion and drift on the retention of memories (Fig. 5A). Because of diffusion, distributions of positions will widen over time, which degrades the ability to distinguish different initial positions of the bump center (Fig. 5A, top). Additionally, directed drift of the dynamics will contract distributions of different initial positions around the same fixed points, making them essentially indistinguishable when read out (Fig. 5A, bottom).

**Fig 5.**
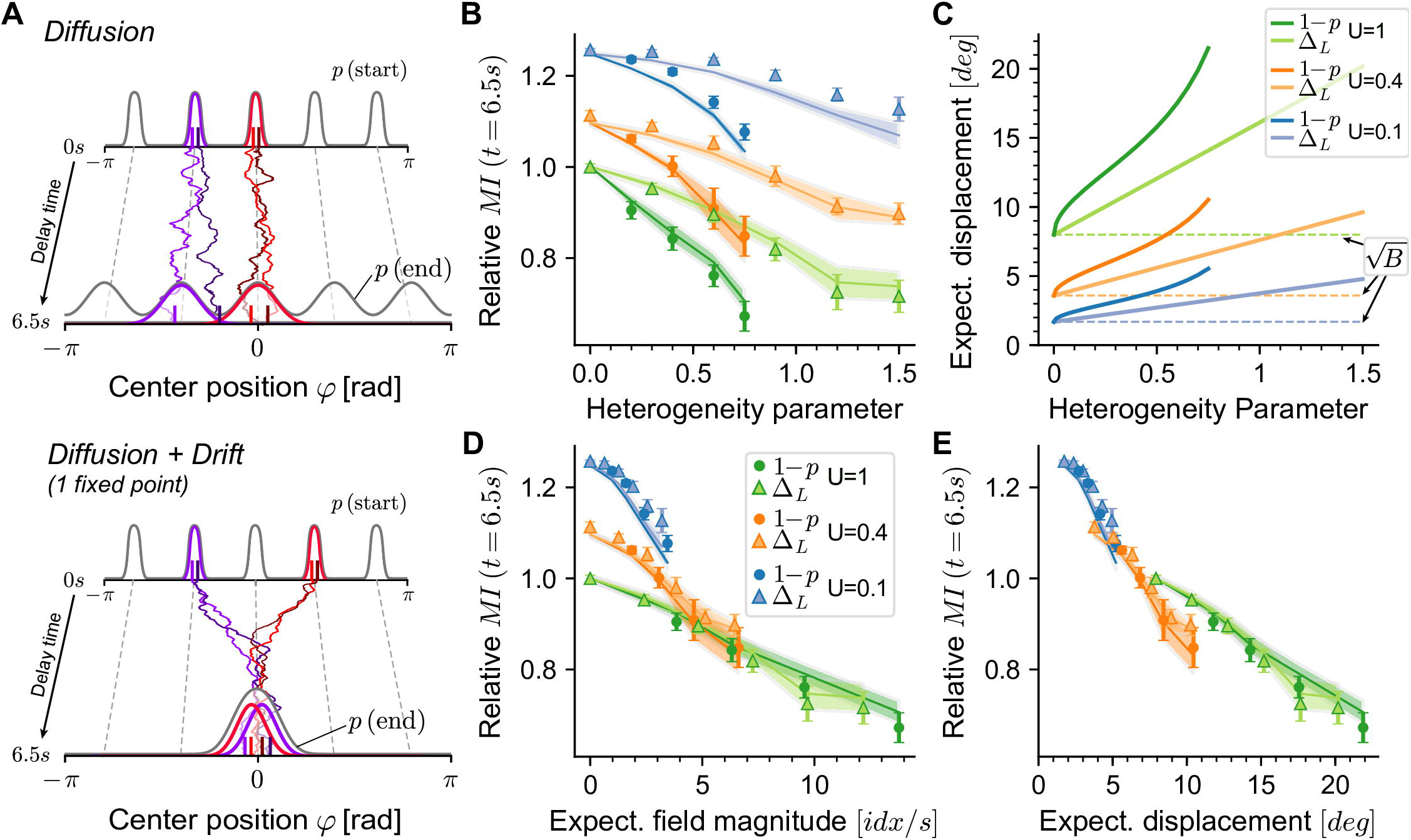
Short-term facilitation increases memory retention. **A** Illustration of the effects of diffusion (top) and additional drift (bottom) on the temporal evolution of distributions of initial positions *p*(start) towards distributions of final positions *p*(end) over 6.5s of delay activity. The bump is always represented by its center position *φ*. Two peaks in the distribution of initial positions *φ*(0) and their corresponding final positions *φ*(6.5) are highlighted by colors (purple, red), together with example trajectories of the center positions. Top: Diffusion symmetrically widens the initial distribution. Bottom: Strong drift towards one single fixed point of bump centers (*φ* = 0) makes the origin of trajectories indistinguishable. **B** Normalized mutual information (MI, see text for details) of distributions of initial and final bump center positions in working memory networks for different STP parameters and heterogeneity parameters(blue: strong facilitation, see legend in panel D). Dots and triangles are average MI (18 – 20 realizations, error bars show 95% CI) obtained from spiking network simulations. Lines show average MI calculated from Langevin dynamics for the same networks, repetitions and realizations (see text, shaded area shows 95% CI). Heterogeneity parameters are *σ_L_* (triangles, in units of *mV*) and 1 − *p* (circles), where *p* is the connection probability. **C** Expected displacement |Δ*φ*| (1*s*) for the same networks as in panel B. Dashed lines indicate displacement induced by diffusion only 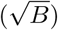, solid lines show the total displacement (including displacement due to drift, calculated as the expected field magnitude 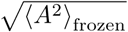). **D** Same as panel B, with x-axis showing the expected field magnitude. **E** Same as panel B, with x-axis showing the expected displacement. In panels B-D, all facilitation parameters except *U* were kept constant at *τ_u_* = 650ms, *τ_x_* = 150ms

As a numerical measure of this ability of such systems to retain memories over the delay period, we turned to mutual information (MI), which provides a measure of the amount of information contained in the readout position about the initially encoded position [55, 56]. To measure MI from simulations (see *Mutual information measure* in Materials and Methods), we analyzed network simulations for varying short-term facilitation parameters (*U*) and magnitudes of frozen noises (*p* and *σ_L_*) (same data set as Fig. 4A,B). We recorded the center positions encoded in the network at the time of cue-offset (*t* = 0) and after 6.5s of delay activity, and used binned histograms (100 bins) to calculate discrete probability distributions of initial (*t* = 0) and final positions (*t* = 6.5). For each trajectory simulated in spiking networks, we then generated a trajectory starting at the same initial position by using the Langevin equation Eq. (4) that describes the drift and diffusion dynamics of center positions. The MI calculated from the resulting distributions of final positions (again at *t* = 6.5) for each network serve as the theoretical prediction for each network. As a reference, we used the spiking network without facilitation (*U* = 1, *τ*_u_ = 650*ms, τ_x_* = 150*ms*) and no frozen noises (*p* =1, *σ_L_* = 0*mV*) and normalized the MI of all other networks (both for spiking simulations and theoretical predictions) with respect to the reference, yielding the measure of *relative MI* presented in Fig. 5B-E.

We found that the relative MI decreased compared to the reference network as network heterogeneities were introduced (Fig. 5B, green). This was expected, since directed drift caused by heterogeneities leads to a loss of information about initial positions. There were two effects of increased short-term facilitation (by decreasing the parameter *U*). First, diffusion was reduced, which was visible in a vertical shift of the relative MI for facilitated networks (Fig. 5A, orange and blue, at 0 heterogeneity).

Second, the effects of frozen noise decreased with increasing facilitation, which was visible in the slopes of the MI decrease (see also S4 Fig). The MI obtained by integration of the Langevin equations (see above) matched those of the simulations well (Fig. 5A, lines). From earlier results, we expected the drift-fields to be slightly over-estimated by the theory as the heterogeneity parameters increase (Fig. 4), which would lead to an under-estimation of MI. We did observe this here, although for *U* = 1 the effect was slightly counter-balanced by the under-estimated level of diffusion (cf. Fig. 3A, right), which we expected to increase the MI. For networks with stronger facilitation (*U* = 0.1), we systematically over-estimated diffusion (cf. Fig. 3, left), and therefore under-estimated MI.

Using our theory, we were able to simplify the functional dependence between MI, short-term plasticity, and frozen noise. Combining the effects of both diffusion and drift into a single quantity for each network, we replaced the field *A*(*φ*) by our theoretical prediction 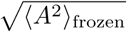 in Eq. (4) and forward integrated the differential equation for a time interval Δ*t* = 1*s*, to arrive at the *expected displacement* in 1*s*:

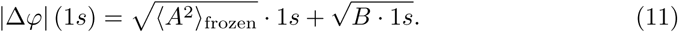

This quantity describes the expected absolute value of displacement of center positions during 1*s*: it increases as a function of the frozen noise distribution parameters (Fig. 5C), but even in the absence of frozen noise it is nonzero due to diffusion. Plotting the MI data in dependence of the first term only 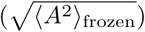, shows that the MI curves collapse onto a single curve for each facilitation parameter (Fig. 5D). Finally, plotting the MI data against |Δ*φ*| (1*s*) we find that all data collapse on to nearly a single curve (Fig. 5E). Thus, the effects of the two sources of frozen noise (corresponding to 〈*A*^2^〉_frozen_) and diffusion (corresponding to *B*) are unified into a single quantity |Δ*φ*| (1*s*).

We performed the same analyses on a large set of network simulations with fixed random connectivity (*p* = 0.6) and varying STP parameters for both depression (*τ_x_*) and facilitation (*U*) (same data set as in Fig. 4C). Increasing the short-term depression time constant *τ_x_* leads to decreased relative MI with a positive offset induced through stronger facilitation (Fig. 6A, blue line). Calculating the expected displacement for these network configurations collapsed the data points mostly onto the same curve as earlier (Fig. 6B). For strong depression combined with weak facilitation (*τ_x_* = 200*ms, U* = 0.8), the drop-off of the relative MI saturates earlier, indicating that for these strongly diffusive networks the effect on MI may not be sufficiently captured by its relationship to |Δ*φ*| (1*s*).

**Fig 6.**
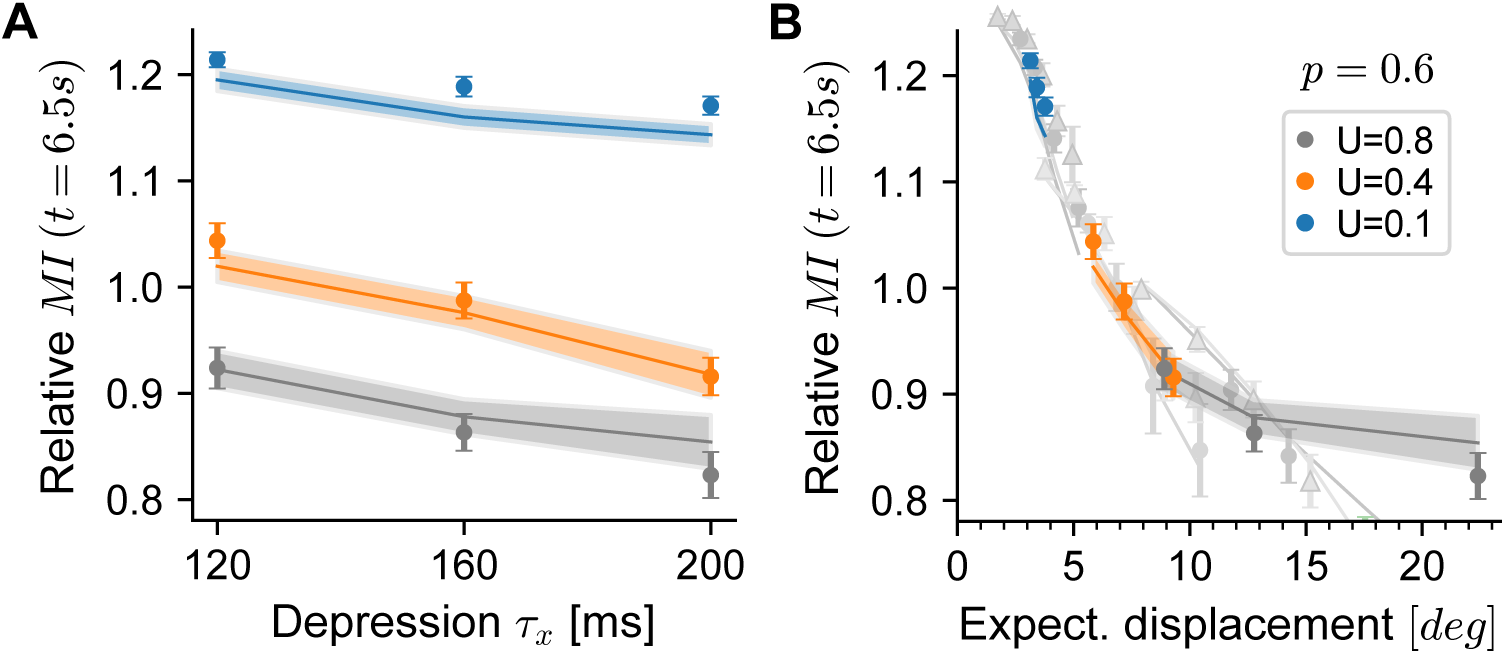
Short-term depression decreases memory retention. **A** Same as Fig. 5B, for network simulations with varying *τ_x_* and *U* (see legend in panel B). MI is normalized to the same value as there. **B** Same as panel A, with x-axis showing the expected displacement. Light gray data points and lines are the data plotted in Fig. 5E. The facilitation time constant was kept constant at *τ_u_* = 650ms.

### Linking theory to experiments: distractors & network size

The large level of abstraction of our theory condenses the complex dynamics of spiking bump attractor networks into a high-level description of a few macroscopic features, which in turn allows matching the theory to behavioral experiments. Here, we demonstrate how such quantitative links could be established using two different features: 1) the sensitivity of the working memory circuit to distractors, and 2) the stability of working memory expressed by the expected displacement. We stress that our model is a simplified description of biological circuits (see Discussion), and thus, at their current level of abstraction, should be seen as a proof of principle rather than a means of deriving accurate quantitative predictions.

#### Predicting the sensitivity to distractor inputs

In a biological setting, drifts introduced by network heterogeneities (frozen noise) could be significantly reduced by (long-term) plasticity [36]. To measure the intrinsic stability of continuous attractor models, earlier studies [11, 46, 57] have alternatively proposed to use *distractor inputs* (Fig. 7A): providing a short external input centered around a position *φ_D_* to the network, the center position of an existing bump state will be biased towards the distracting input, with stronger biases appearing for closer distractors. Distractors forcibly expose the working memory system’s intrinsic time scale by creating externally induced drifts, thereby testing its sensitivity (in terms of distractor distance), and they are equally implementable in behavioral experiments [57].

**Fig 7.**
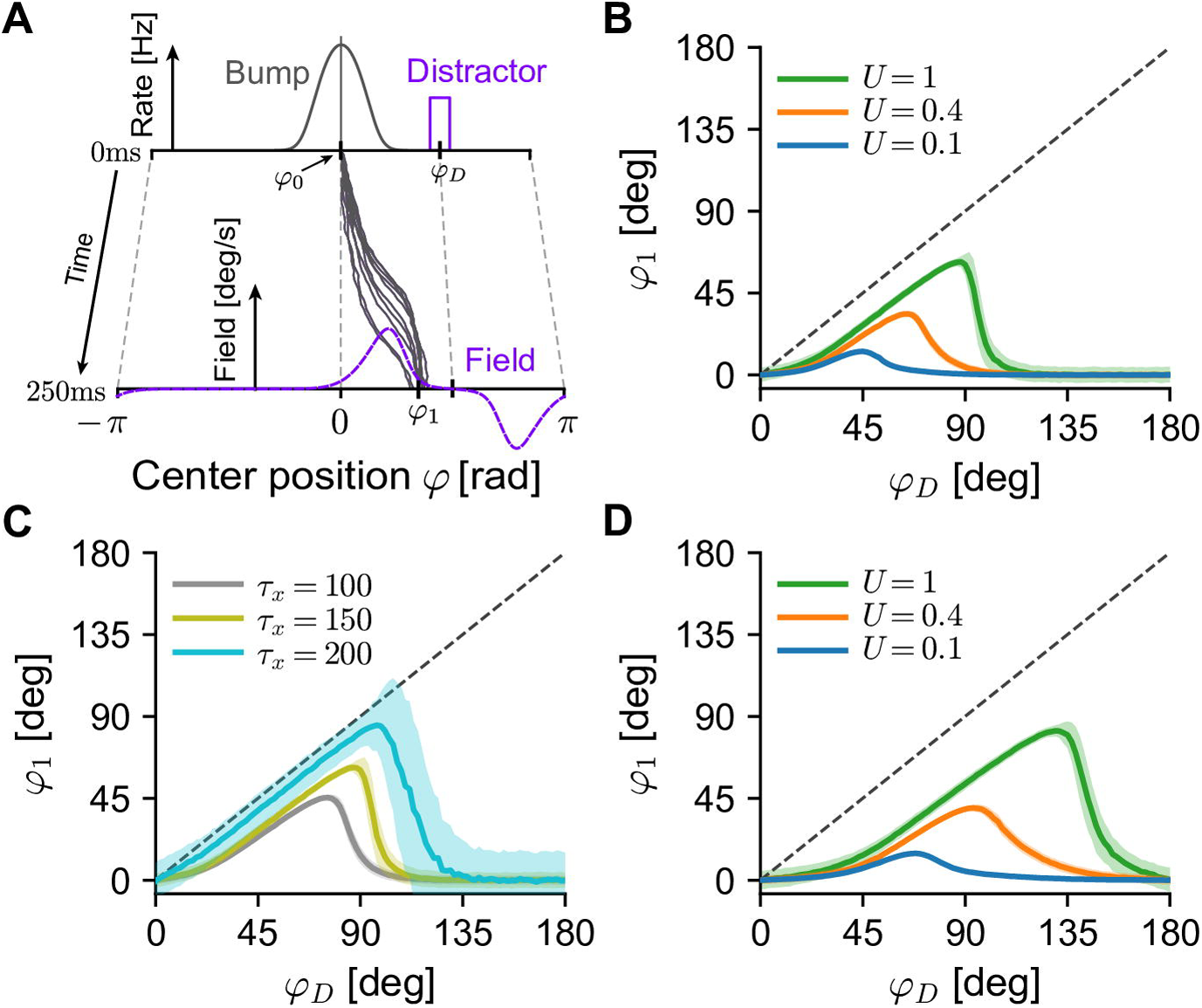
Effect of short-term plasticity on distractor inputs. **A** While a bump (“Bump”) is centered at an initial angle *φ*_0_ (chosen to be 0), additional external input causes neurons centered around the position *φ*_*D*_ to fire at elevated rates (“Distractor”). The theory predicts the shape and magnitude of the induced drift field (“Field”) and the mean bump center *φ*_1_ after 250ms of distractor input. Gray trajectories are example simulations of bump centers of the corresponding Langevin equation Eq. (4). **B** Mean final positions *φ*_1_ of bump centers (1000 repetitions, shaded areas show 1 standard deviation) as a function of the distractor input location *φ*_*D*_. Increased short-term facilitation (blue: strong facilitation, *U* = 0.1; orange: intermediate facilitation, *U* = 0.4; green: no facilitation *U* = 1) leads to less displacement due to the distractor input. Other STP parameters were kept constant at *τ_u_* = 650ms, *τ_x_* = 150ms. **C** Same as panel B, for three different depression time constants *τ_x_*, while keeping *U* = 0.8, *τ_u_* = 650ms fixed. **D** Same as panel B, with a broader bump half-width (*σ_g_* = 0.8rad ≈ 45.8deg). All other panels use the same bump half-width as in the rest of the study (*σ_g_* = 0.5rad ≈ 28.7deg) (see S1 Fig).

Our theory can readily yield quantitative predictions for the distractor paradigm. To accommodate distractor inputs in the theory, we assume that they cause some units *i* to fire at elevated rates *ϕ*_0,*i*_ + Δ*ϕ*_*i*_, which will introduce a drift field according to Eq. (7) (Fig. 7A, purple dashed line). The resulting dynamics (Eq. (4)) of diffusion and drift during the presentation of the distractor input then allow us to calculate the expected shift of center positions as a function of all network parameters, including those of short-term plasticity. Repeating this paradigm for varying positions of the distractor inputs (see *Distractor analysis* in Materials and Methods for details), our theory predicts that strong facilitation will strongly decrease both the effect and radial reach of distractor inputs (Fig. 7B, blue), when compared to the unfacilitated system (Fig. 7B, green) – in qualitative agreement with simulation results involving a related (cell-intrinsic) stabilization mechanism [46]. Conversely, we predict that longer recovery from short-term depression tends to increase the sensitivity to distractors (Fig. 7C). The displacement caused by each distractor input is given integrating the resulting dynamics of Eq. (4) over the stimulus duration. As such, the magnitude of the displacement will increase both with the amplitude and the duration of the distractor input. Finally, our theory demonstrates that the bump shape, importantly bumps with a broader width, can also significantly increase the effect and radial reach of distractor inputs (Fig. 7D).

#### Constraining the size of working memory networks

The simple theoretical measure of expected displacement |Δ*φ*| (1*s*) introduced in the last section can be related to behavioral experiments: a value of |Δ*φ*| (1*s*) = 1.0 deg lies in the upper range of experimentally reported deviations due to diffusive and systematic errors in behavioral studies [58, 59]. What are the microscopic circuit compositions that can attain such a (high) level of working memory stability? In particular, since an increase in network size can reduce diffusion [11] and the effects of random heterogeneities [16, 36, 38, 45], we turned to the question: *which networks sizes would be needed to yield this level of stability in a one-dimensional continuous memory system*?

To address the question of network size, we extended our theory to include the size *N* of the excitatory population as an explicit parameter (see *System size scaling* in Materials and Methods for details). We find (see Eq. (59)) that the expected field magnitude 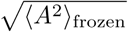 (cf. Eq. (10)) scales with the connectivity parameter *p* and the system size *N* to leading order as 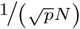, whereas the perturbations due to heterogeneity of leak potentials scale as 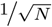, both in accordance with earlier results [16, 36, 38, 45]. For the diffusion scale 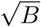 (cf. Eq. (5)) we find a scaling as as 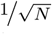, also in agreement with earlier work [11, 14, 36, 39, 45]. Using numerical coefficients in Eq. (4) extracted from the spiking simulation of a reference network (*U* = 1, *τ_x_* = 150 and *N_e_* = 800), we extrapolated the theory by changing the system size *N* and short-term plasticity parameters. We then constrained parameters of our theory by published data (Table 1). Short-term plasticity parameters were based on two groups of strongly facilitating synapses found in a study of mammalian (ferret) prefrontal cortex [60]. The same study reported a general probability *p* = 0.12 of pyramidal cells to be connected. However, for pairs of pyramidal cells that were connected by facilitating synapses, the study found a high probability of reciprocal connections (*p_rec_* = 0.44): thus if neuron A was connected to neuron B (with probability *p*), neuron B was connected to neuron A with high probability (*p_rec_*), resulting in a non-random connectivity. To approximate this in the random connectivities supported by our theory, we evaluated a second, slightly elevated, level of random connectivity, that has the same mean connection probability as the non-random connectivity with these additional reciprocal connections: *p* + *p* · *p_rec_* = 0.1728. For the standard deviation of leak reversal-potentials *σ_L_*, we used values measured in two studies [61, 62].

**Table 1.**
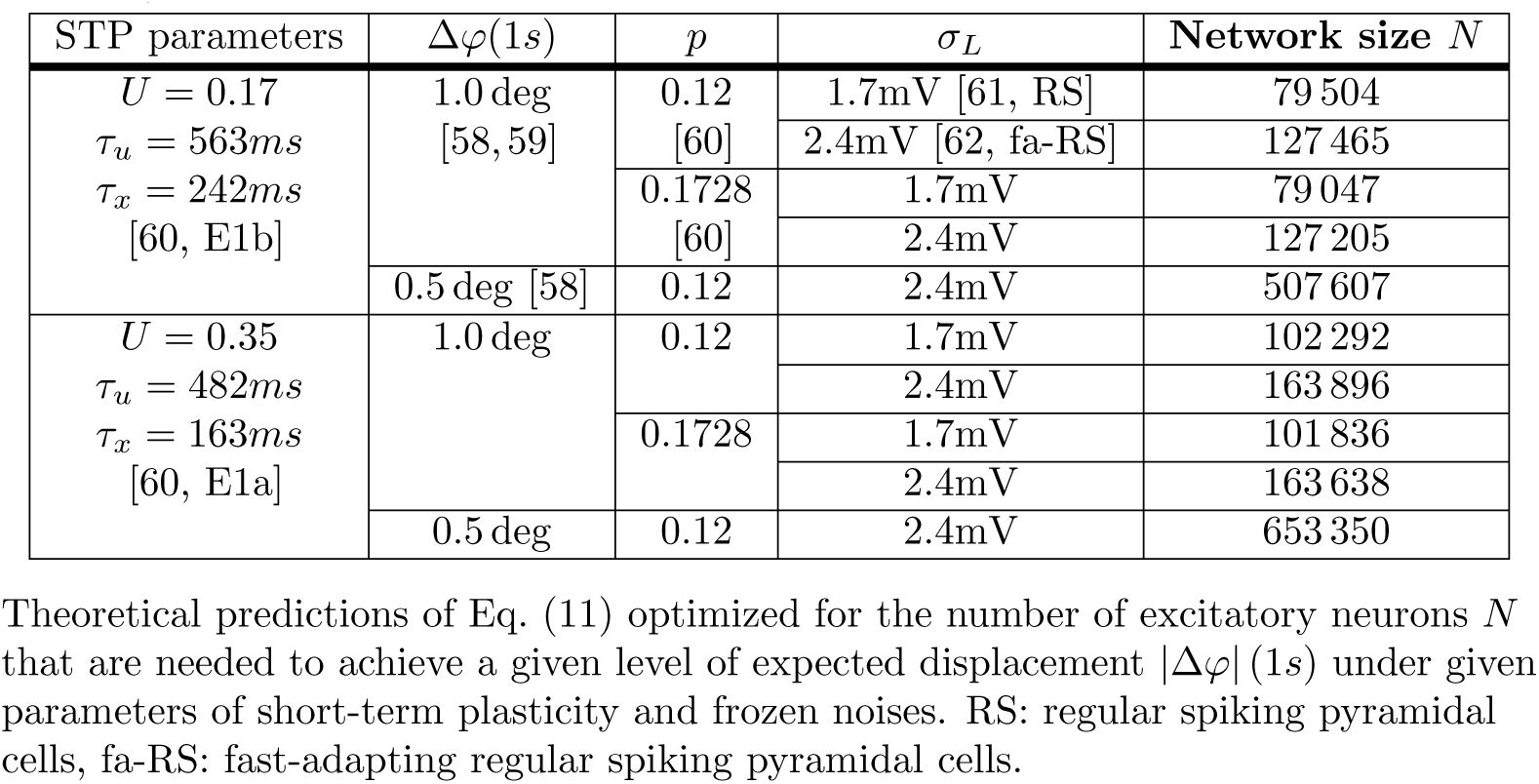
Upper bounds on system-sizes for stable continuous attractor memory in prefrontal cortex.

The resulting theory makes quantitative predictions for combinations of network size *N* and all other parameters that yield the desired levels of working memory stability (Table 1, see also S5 Fig). Network sizes were all smaller than 10^6^ neurons, with values depending most strongly on the value of the facilitation parameter *U* and the magnitude of the leak reversal-potential heterogeneities *σ_L_*. Since the expected field magnitude scales weakly 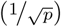 with the recurrent connectivity *p*, increasing *p* lead only to comparatively small decreases in the predicted network sizes. Finally, we see that the increasing the reliability of networks comes at a high cost: decreasing the expected displacement to |Δ*φ*| (1*s*) = 0.5 deg [58] increases the required number of neurons by nearly a number of 4 for both facilitation settings we investigated. Nevertheless, these network sizes still lie within anatomically reasonable ranges [63]. In summary, our theory predicts that, given the simplifying assumptions of our models, continuous attractor networks with realistic values for the strength of facilitation and depression of recurrent connections could achieve sufficient stability, even in the presence of a realistic degree biological variability.

## Discussion

We presented a theory of drift and diffusion in continuous working memory models, exemplified on a one-dimensional ring attractor model. Our framework extends earlier approaches calculating the effects of noise by projection onto the attractor manifold [37, 39, 40] by including the effects of short-term plasticity. Our approach further generalizes earlier work on drift in continuous attractors with short-term plasticity [38] to include diffusion and the dynamics of short-term depression. The theory predicts that facilitation and depression play opposite roles in making continuous attractors robust against the influences of both dynamic noise (introduced by spiking variability) and frozen noise (introduced by biological variability). We have confirmed the quantitative predictions of our theory in simulations of a ring-attractor implemented in a spiking network model with synaptic facilitation and depression, and found theory and simulation to be quantitatively in good agreement.

In *Short-term plasticity controls memory retention*, we demonstrated the effects that STP has through drift and diffusion on the information retained in continuous working memory. Using our theoretical predictions of drift and diffusion we were able to derive the expected displacement |Δ*φ*| as a function of STP parameters and the frozen noise parameters, which provides a simple link between the resulting Langevin dynamics of bump centers and mutual information (MI) as a measure of working memory retention. Our results can be generalized in several directions. First, the choice of 1s of forward integrated time for |Δ*φ*| (Eq. (11)) was arbitrary. While a choice of ~ 2*s* lets the curves in Fig. 5E collapse slightly better, we chose 1s to avoid further heuristics. Second, we expect values of MI to decrease as the length of the delay period is increased. Our choice of 6.5*s* is comparable to delay periods often considered in behavioral experiments (usually 3-6s) [59, 64, 65]. However, a more rigorous link between the MI measure and the underlying attractor dynamics would be desirable. Indeed, for noisy channels governed by Fokker-Planck equations, this might be feasible [66], but goes beyond the scope of this work.

In *Linking theory to experiments: distractors & network size*, we demonstrated that the high-level description of the microscopic dynamics obtained by our theory allows its parameters to be constrained by experiments. Considering that our model is a simplified description of its biological counterparts (see next paragraph), these demonstrations are to be seen as a proof of principle, which will not be able to yield biologically accurate predictions in its current level of abstraction. However, since distractor inputs can be implemented in silico as well as in behavioral experiments (see e.g. [57]), they could eventually provide a quantitative link between the continuous attractor models and working memory systems, by matching the resulting distraction curves. Given the simplifying assumptions, our theory provides a direct map between underlying network parameters and their combined effects on distraction sensitivity, facilitating previous approaches in which these distraction curves had to be extracted through repeated microscopic simulations for single parameter settings [46]. We further used our theory to derive bounds on network parameters, in particular the size of networks, that lead to “tolerable” levels of drift and diffusion in the simplified model. For large magnitudes of frozen noise our theory tends to over-estimate the expected magnitude of drift-fields slightly (cf. Fig. 4). Thus, we expect the predictions made here to be upper bounds on network parameters needed to achieve a certain expected displacement. Finally, while the predictions of our theory might deviate from biological networks, they could be applied to accurately characterize the stability of, and the effects of inputs to, bump attractor networks implemented in neuromorphic hardware for robotics applications [67].

Our results show, to our knowledge for the first time, that strong facilitation (small values of *U*) does not only slow down directed drift [38], but also efficiently suppresses diffusion in continuous attractor models. However, in delayed response tasks involving saccades, that presumably involve continuous attractors in the prefrontal cortex [11, 22], one does observe an increase of variability in time [64]: both quickly accumulating systematic errors (alike drift) [59] and more slowly increasing variable errors (with variability growing linear in time, alike diffusion) [58] appear. Indeed, there are several other possible sources of variability in cortical working memory circuits, which we did not consider here. For example, noisy synaptic transmission and STP [68] or heterogeneous STP parameters as found in [60]. These will probably induce further drift and diffusion and the effectiveness of facilitation as a stabilizing mechanism in coping with them remains to be investigated. For simplicity, we also excluded AMPA currents from the recurrent excitatory interactions, which are found in biological neuronal networks [11]. However, since STP acts by presynaptic scaling of neurotransmitter release, it will act symmetrically on both AMPA and NMDA receptors, which we thus expect to lead to similar dynamical effects of STP as those reported here and which might be amenable to a similar analytical approach. Additionally, variable errors might be introduced elsewhere in the pathway between visual input and motor output (but see [69]) or by input from of other noisy local circuits during the delay period [70].

Several additional dynamical mechanisms might also influence the stability of continuous attractor working memory circuits. For example, intrinsic neuronal currents that modulate the neuronal excitability [46] or firing-rate adaptation [71] have been shown to affect bump stability. It should be noted that many such dynamical processes could be accommodated in our theoretical approach, by including their linearized dynamics in the calculation of the projection vector (cf. *Projection of dynamics onto the attractor manifold* in Materials and Methods). Fast corrective inhibitory feedback has also been shown to stabilize spatial working memory systems (in balanced networks) [72]. It has been shown that, on the timescale of hours to days, homeostatic processes can be used to counteract the drift introduced by frozen noise [36]. Finally, inhibitory connections that are distance-dependent [11] and show short-term plasticity [73] could also influence bump dynamics.

We have focused here on ring-attractor models that obtain their stable firing-rate profile due to perfectly symmetric connectivity. Our approach can also be employed to analyze ring-attractor networks with short-term plasticity, in which weights show (deterministic or stochastic) deviations from symmetry (see *Frozen noise* in Materials and Methods for stochastic deviations). Although not investigated here, continuous line-attractors arising through a different weight-symmetry should be amenable to similar analyses [39]. Finally, it should be noted that adequate structuring of the recurrent connectivity can also positively affect the stability of continuous attractors [14]. For example, translational asymmetries included in the structured heterogeneity can break the continuous attractor into several isolated fixed points, which can lead to decreased diffusion along the attractor [56].

We provided evidence that short-term synaptic plasticity controls the sensitivity of attractor networks to both fast diffusive and frozen noise. Control of short-term plasticity via neuromodulation [74] would thus represent an efficient “crank” for adapting the timescale of computations of such networks. By changing the properties of presynaptic calcium entry [75], inhibitory modulation mediated via GABA_B_ and adenosine A_1_ receptors can lead to increased facilitatory components in rodent cerebellar [76] and avian auditory synapses [77]. Dopamine, serotonin and noradrenaline have all been shown to differentially modulate short-term depression (and facilitation when blocking GABA receptors) at sensorimotor synapses [78]. Interestingly, next to short-term facilitation on the timescale of seconds, other dynamic processes up-regulate recurrent excitatory synaptic connections in prefrontal cortex [60]: synaptic augmentation and post-tetanic potentiation operate on longer time scales (up to tens of seconds), and might be able to support working memory function [79]. While the long time scales of these processes might again render putative short-term memory networks inflexible, there is evidence that they might also be under tight neuromodulatory control [80]. Neuromodulation could also generally provide higher flexibility (see above) to a facilitation-stabilized working memory system, by allowing for more dynamical control of the rigidity of memory representations [46].

### Comparison to earlier work

Similar to an earlier theoretical approach using a simplified rate model [38], we find that the slowing of drift by facilitation depends mainly on the facilitation parameter *U*, while the time constant *τ_u_* has a less pronounced effect. While the approach of [38] relied on the projection of frozen noise onto the derivative of the first spatial Fourier mode of the bump shape along the ring, here we reproduce and extend this result (1) for arbitrary neuronal input-output relations and (2) a more detailed spatial projection that involves the synaptic dynamics and the bump shape. While, our theory can also accommodate noisy recurrent connection weights as frozen noise, as used in in [38] (see *Frozen noise* in Materials and Methods for derivations), the drifts generated by these heterogeneities were generally small compared to diffusion and the other sources of heterogeneity.

A second study investigated short-term facilitation and showed that it reduces drift and diffusion in a spiking network, for a fixed setting of *U* (although the model of short-term facilitation differs slightly from the one employed here) [46]. Contrary to what we find here, these authors find that an increase in *τ_u_* leads to increased diffusion, while we find that an increase over the range they investigated (~ 0.5*s* – 4*s*) would decrease the diffusion by a factor of nearly two. More precisely, for our shape of the bump state (which we keep fixed) we predict a reduction from ~ 26 to ~ 16 *deg*^2^/*s* for a similar setting of facilitation *U*. These differences might arise from an increasing width of the bump attractor profile for growing facilitation time constants in [46], which would then lead to increased diffusion in our model. Whether this effect persists under the two-equation model of saturating NMDA synapses used there remains to be investigated. Finally, increasing the time constant of recurrent NMDA conductances has been shown to also reduce diffusion [46], in agreement with our theory, according to which the normalization constant *S* increases with *τ_s_* [39].

In a study that investigated only a single parameter value for depression (*τ_x_* = 160*ms*, no facilitation) in a spiking network similar to the one investigated here [44], the authors observed no apparent effect of short-term depression on the stability of the bump. In contrast, we find that stronger short-term depression will indeed increase both diffusion and directed drift along the attractor. Our result agrees qualitatively with earlier studies in rate models, which showed that synaptic depression, similar to neuronal adaptation [10, 81], can induce movement of bump attractors [42, 43, 82, 83]. In particular, the study of [42] showed that for simpler rate models a regime exists where the bump state moves with constant speed along the attractor manifold. We did not find any such directed movement in our networks, which could be due to fast spiking noise stabilizing the directed movement of bumps observed in noise-free rate models, as shown in [81]: there, additive fast noise was able to cancel directed bump movement caused by single neuron adaptation.

### Extensions & Shortcomings

The coefficients of Eq. (4) give clear predictions as to how drift and diffusion will depend on the shape of the bump state and the neural transfer function *F*. The relation is not trivial, since the pre-factors *C_i_* and the normalization constant *S* also depend on the bump shape. For the diffusion strength Eq. (5), we explored this relation numerically, by artificially varying the shape of the firing rate profile (while extrapolating other quantities). Although a more thorough analysis remains to be performed, a preliminary analysis shows (see S6 Fig) that diffusion increases both with bump width and top firing rate, consistent with earlier findings [11, 32].

In *Linking theory to experiments: distractors & network size*, we have demonstrated that our theory can be used to predict the shape and effect of drift fields that are generated by localized external inputs due to distractor inputs. More generally, any localized external input (excitatory or inhibitory) will cause a deviation Δ*ϕ*_i_ from the steady-state firing rates, which, in turn, generates a drift field by Eq. (7). This could predict the strength and location of external inputs that are needed to induce continuous shifts of the bump center at given speeds, for example when these attractor networks are designed to track external inputs (see e.g. [10, 84]). It should be noted that in our simple approximation of this distractor scheme, we assume the system to remain at approximately steady-state, i.e. that the bump shape is unaffected by the additional external input, except for a shift of the center position. For example, we expect additional feedback inhibition (through the increased firing of excitatory neurons caused by the distractor input) to decrease bump firing rates. A more in depth study and comparison to simulations will be left for further work.

The spiking networks we analyzed here are tuned to display balanced inhibition and excitation in the inhibition dominated uniform state [85, 86], while the bump state relies on positive currents, mediated through strong recurrent excitatory connections (cf. [44] for an analysis). Similar to other spiking network models of this class, this mean-driven bump state shows relatively low variability of neuronal inter-spike-intervals, because near the bump center the mean input is close to (or even above) the firing threshold [87, 88] (see also next paragraph). While the decreased variability appears for neurons in the center of the firing rate profile, we found that neurons at its flanks still display variable firing, with statistics close to that expected of spike trains with Poisson statistics (see S7 Fig), which may be because the flank’s position slightly jitters. Since the non-zero contributions to the diffusion strength are constrained to these flanks (cf. Fig. 1D), the simple theoretical assumption of Poisson statistics of neuronal firing still matches the spiking network quite well. As discussed in *Short-term plasticity controls diffusion*, we find that our theory over-estimates the diffusion as bump movement slows down for small values of *U* – this may be due to a decrease in firing irregularity in stable bumps, at which the Poisson assumption becomes inaccurate.

More recent bump attractor approaches allow networks to perform working memory function with a high firing variability also during the delay period [3], in better agreement with experimental evidence [89]. These networks show bi-stability, where both stable states show balanced excitation and inhibition [87] and the higher self-sustained activity in the delay activity is evoked by an increase in fluctuations of the input currents (noise-driven) rather than an increase in the mean input [90]. This was also reported for a ring-attractor network (with distance-dependent connections between all populations), where facilitation and depression are crucial for irregularity of neuronal activity in the self-sustained state [45]. Application of our approach to these setups is left for future work.

## Materials and methods

### Analysis of drift & diffusion with STP

For the following, we define a concatenated 3 · *N* dimensional column vector of state variables **y** = (**s**^*T*^, **u**^*T*^, **x**^*T*^)^*T*^ of the system Eq. (3). Given a (numerical) solution of the stable firing rate profile 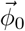 we can calculate the stable fixed point of this system by setting the l.h.s. of Eq. (3) to zero. This yields steady-state solutions for the synaptic activations, facilitation and depression variables **y**_0_ = (**s**_0_, **u**_0_, **x**_0_):

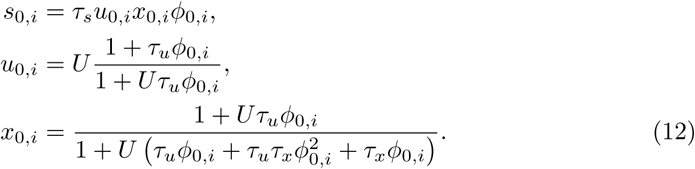

We then linearize the system Eq. (3) at the fixed point **y**_0_, introducing a change of variables consisting of perturbations around the fixed point: **y** = **y**_0_ + *δ***y** = **y**_0_ + (*δ***s**^*T*^, *δ***u**^*T*^, *δ***x**^*T*^) and *ϕ*_*i*_ = *ϕ*_0,*i*_ + *δϕ_i_*. To reach a self-consistent linear system, we further assume a separation of time scales between the neuronal dynamics and the synaptic variables, in that the neuronal firing rate changes as an immediate function of the (slow) input. This allows replacing 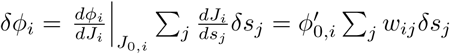, where we introduce the shorthand 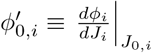. Finally, keeping only linear orders in all perturbations, we arrive at the linearized system equivalent of Eq. (3):

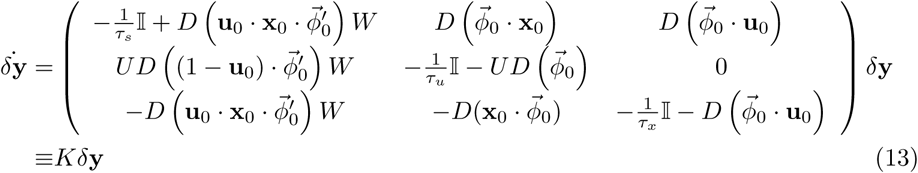

Here, dots between vectors indicate element-wise multiplication, the operator *D*: ℝ^*n*^ → ℝ^*n*×*n*^ creates diagonal matrices from vectors, and *W* = (*w_ij_*) is the synaptic weight matrix of the network.

#### Projection of dynamics onto the attractor manifold

To project the dynamical system Eq. (13) onto movement of the center position *φ* of the firing rate profile, we assume that *N* is large enough to treat the center position *φ* as a continuous variable. We also assume that the network implements a ring-attractor: the system dynamics are such that the firing rate profile 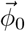 can be freely shifted to different positions along the ring, changing the center position *φ*, while retaining the same shape. All other possible directions of change in this system are assumed to be constrained by the system dynamics. In the system at hand, this implies that the matrix *K* of Eq. (13), which captures the linearized dynamics around any of these fixed points, will have a *zero eigenvalue* corresponding to the eigenvector of a change of the dynamical variables under a change of position *φ*, while all other eigenvalues are negative [39].

Formally, the column eigenvector to the eigenvalue 0 is given by changes in the state variables as the bump center position *φ* is translated along the manifold:

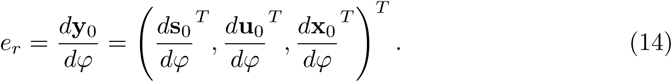

Let *e_l_* be the associated row left-eigenvector (also to eigenvalue 0) of *K*, normalized such that:

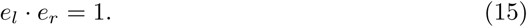

In Section 1 of S1 Supporting Information, we show that the eigenvector *e_l_* projects the system Eq. (13) onto dynamics of of the center position:

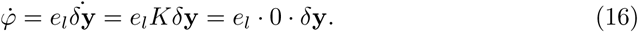

Under the linearized ring-attractor dynamics *K*, the center position is thus not subject to any dynamics, making it susceptible to any displacements by noise.

#### Calculation of the left eigenvector *e_l_*

If the matrix *K* is symmetric, the left and right eigenvectors *e_l_* and *e_r_* for the same eigenvalue 0 are the transpose of each other. Unfortunately, here this is not the case (see Eq. (13)), and we need to compute the unknown vector *e_l_*, which will depend on the coefficients of the known vector *e_r_*. In particular, we look for a parametrized vector **y**′(**y**) = (**t**^*T*^(**y**), **v**^*T*^(**y**), **z**^*T*^(**y**))^*T*^ that for **y** = *e_r_* fulfills the transposed eigenvalue equation of the left eigenvector:

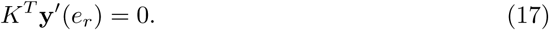

In Section 2 of S1 Supporting Information, we derive variables **y**′ that fulfill the transposed dynamics **ẏ**′ = *K^*T*^***y**′ and for which it holds that **ẏ**′(*e_r_*) = 0, thus fulfilling the condition Eq. (17). In this case we know that (due to uniqueness of the 1-dimensional eigenspace associated to the 0 eigenvalue) the vector **y**′^*T*^ is proportional to *e_l_*:

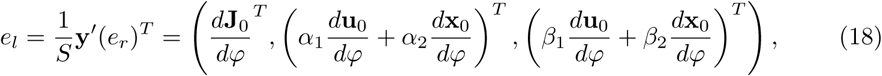

where *S* is a proportionality constant and 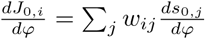 is the change of the steady-state input arriving at neuron *i* under shifts of the center position *φ*.

Finally, the proportionality constant *S* can be calculated by using Eq. (18) in Eq. (15) (see Section 3 of S1 Supporting Information for details):

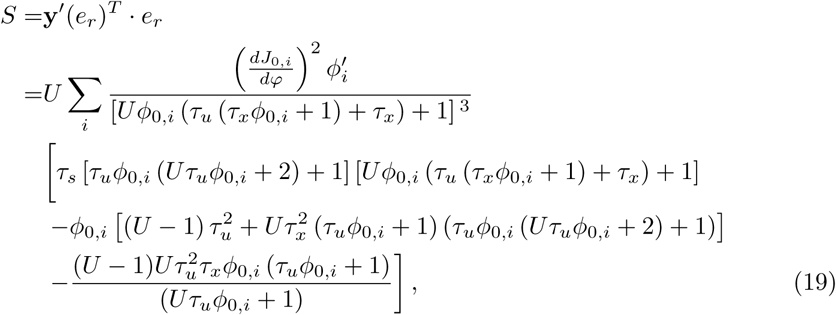

where 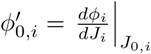 is the linear change of the firing rate of neuron *i* at its steady-state input *J*_0,*i*_.

#### Diffusion

To be able to describe diffusion on the continuous attractor, we need to extend the model by a treatment of the noise induced into the system through the variable process of neuronal spike emission. Starting from Eq. (3), we assume that neurons *i* fire according to independent Poisson processes 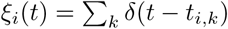, where *t_i,k_* is a Poisson point process with time-dependent rate *ϕ_i_*, The variability of the point process *ξ_i_*(*t*) introduces noise in the synaptic variables. By neglecting the shot-noise (jump-like) nature of this process, we can capture the neurally induced variability simply as white noise with variance proportional to the incoming firing rates [47], 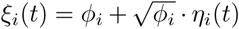, where *η_i_* are white Gaussian noise processes with mean 〈*η_i_*〉 = 0, and correlation function 〈*η_i_*(*t*)*η_j_*(*t*′)〉 = *δ*(*t* − *t*′)*δ_ij_*. This model of *ξ_i_*(*t*) preserves the mean and the auto-correlation function of the original Poisson processes. Here, we introduce diffusive noise for each synaptic variable separately, but later average their *linear* contributions over the large population, when projecting onto movement along the continuous manifold (see below, and also [39, Supplementary Material] for a discussion).

Substituting the noisy processes *ξ_i_*(*t*) for *ϕ_i_*(*t*) in Eq. (3) results in the following system of 3 · *N* coupled Ito-SDEs:

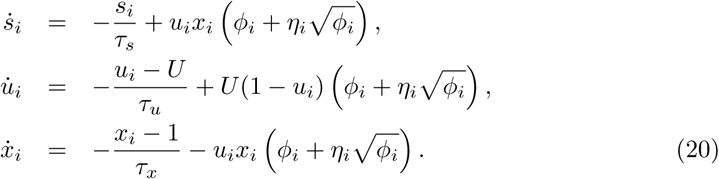

Note that the noise inputs *η_i_* to the synaptic variables for neuron *i* are all identical, since they result from the same presynaptic spike train.

Linearizing this system around the noise-free steady-state Eq. (12) and considering only the unperturbed noise (we neglect multiplicative noise terms by replacing the terms 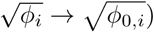), we arrive at the linearized system equivalent of Eq. (20):

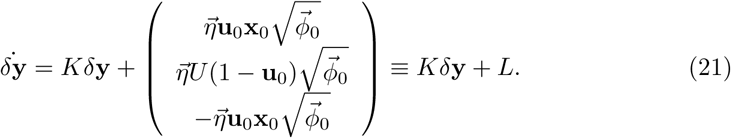

Note that the same vector of white noises 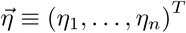 appears three times.

Left-multiplying this system with the eigenvector e_l_ yields a stochastic differential equation for the center position (cf. Eq. (16)):

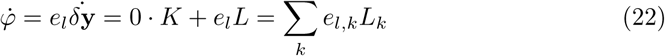

Through the normalization by *S* (Eq. (18)), which sums over all neurons, the individual contributions *e_l,k_* become small as the number of neurons *N* increases (this scaling is made explicit in *System size scaling*). Thus, for large networks we average the small contributions of many single noise sources, which validates the diffusion approximation above.

In Section 4 of S1 Supporting Information, we show that we can rewrite Eq. (22) by introducing a single Gaussian white noise process with intensity *B* (Eq. (5) of the main text), that matches the correlation function of the summed noises:

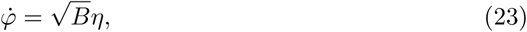

where *η* is a white Gaussian noise process with 〈*η*〉 = 0 and 〈*η*(*t*)*η*(*t*′)〉 = *δ*(*t* − *t*′). Note, that the value of *B* is the same under changes of the center position *φ*: these correspond to index-shifting (mod *N*) all vectors in Eq. (5), which leaves the sum invariant.

#### Drift

While the linearization and diffusion coefficient calculated above are invariant with respect to shifts of the bump center, the directed drift introduced by frozen variability will depend on the bump center position *φ*. Let the center position of the bump be 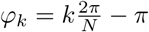 (for 0 ≤ *k* < *N*) with the corresponding firing rate profile 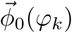, which is an index-shifted version of the *ϕ*_0,*i*_(*φ_k_*) = *ϕ*_0,*i*−*k*_(*φ*_0_).

To calculate the directed drift for a given position *φ_k_*, we first rotate the system such that the steady-state firing rate profile is again centered at *φ*_0_ = −*π*. Then, we assume that the effect of all sources of frozen variability can be expressed by a vector of small firing rate perturbations 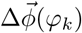 to the steady-state firing rate profile:

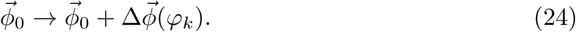

These firing rate perturbations stem from any deviation of the neural system from the “baseline” case and change with each center position *φ_k_* of the bump. The resulting drift field will thus depend on the center position around which we linearize the synaptic dynamics in the following. In *Frozen noise* we will calculate the perturbations induced by random network connectivity, as well as heterogeneous leak reversal-potentials in excitatory neurons of the spiking network.

The firing rate perturbations Eq. (24) add an additional term in the linearized equations Eq. (21):

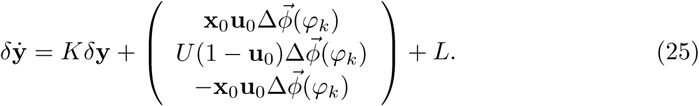

As before, we left-multiply by the left eigenvector *e_l_*, thereby projecting the dynamics onto changes of the center position. This eliminates the linear response kernel *K* and yields a drift-term in the formerly purely diffusive SDE Eq. (23) (see Section 5 of S1 Supporting Information for details):

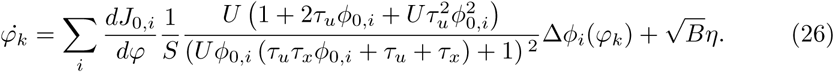

We again assume that the number of neurons *N* is large enough to treat the center position as a continuous variable *φ* ∈ [−*π,π*), allowing us to treat the set of point-wise linearizations of the dynamics at *φ_k_* as a drift-field *A*(*φ*) in Eq. (7) of the main text.

For this, it is important to note that, even for random sources of variability, the first term in the point-wise linearization Eq. (26) at each *φ_k_* will vary nearly continuously with changes in the center position. Intuitively, the sum weighs the vector 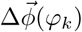 of random numbers with a smooth function of the smoothly varying firing-rate profile 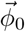. Shifts in the center position *φ_k_* yield (to first order) index-shifts in the vector of random numbers (see *Frozen noise*), equivalent to index-shifts of the firing-rate profile. Thus, small changes in center positions will lead to small changes in the summands of Eq. (26). While our results validate the approach, a more rigorous proof of these arguments will be left for future work.

### Spiking network model

Spiking simulations are based on a variation of a popular ring-attractor model of visuospatial working memory of [11] (and used with variations in [27, 29, 32, 36, 46]). The recurrent excitatory connections of the original network model have been simplified, to allow for faster simulation as well as analytical derivations of the recurrent synaptic activation. The implementation details are given below, however the major changes are: 1) all recurrent excitatory conductances are voltage independent; 2) a model of synaptic short-term plasticity via facilitation and depression [48, 91, 92] is used to dynamically regulate the weights of the incoming spike-trains 3) recurrent excitatory conductances are computed as linear filters of the weighted incoming spike trains instead of the second-order kinetics for NMDA saturation used in [11].

#### Neuron model

Neurons are modeled by leaky integrate-and-fire dynamics with conductance based synaptic transmission [11, 49]. The network consists of recurrently connected populations of *N_E_* excitatory and *N_I_* inhibitory neurons, both additionally receiving external spiking input with spike times generated by *N*_ext_ independent, homogeneous Poisson processes, with rates *v_ext_*. We assume that external excitatory inputs are mediated by fast AMPA receptors, while, for simplicity, recurrent excitatory currents are mediated only by slower NMDA channels.

The dynamics of neurons in both excitatory and inhibitory populations are governed by the following system of differential equations indexed by *i* ∈ {0,…, *N_E/I_* − 1}:

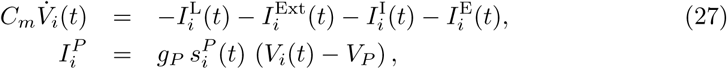

where *P* ∈ {L,Ext,I,E}, *V* denotes voltages (membrane potential) and *I* denotes currents. Here, *C*_m_ is the membrane capacitance and *V*_L_, *V*_E_, *V*_I_ are the reversal potentials for leak, excitatory currents, and inhibitory currents, respectively. The parameters *g_P_* for *P* ∈ {L,Ext,I,E} are fixed scales for leak (L), external input (Ext) and recurrent excitatory (E) and inhibitory (I) synaptic conductances, which are dynamically gated by the unit-less gating variables 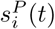. These gating variables are described in detail below, however we set the leak conductance gating variable to 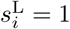. For excitatory neurons, we refer to the excitatory and inhibitory conductance scales by *g*_EE_ ≡ *g*_E_ and *g*_EI_ ≡ *g*_I_, respectively. Similarly, for inhibitory neurons, we refer to the excitatory and inhibitory conductance scales by *g*_IE_ ≡ *g*_E_ and *g*_II_ ≡ *g*_I_, respectively.

The model neuron dynamics (Eq. 27) are integrated until their voltage reaches a threshold *V*_thr_. At any such time, the respective neuron emits a spike and its membrane potential is reset to the value *V*_res_. After each spike, voltages are clamped to *V*_res_ for a refractory period of *τ*_ref_. See the Tables in S1 Table and S2 Table for parameter values used in simulations.

#### Synaptic gating variables and short-term plasticity

The unit-less synaptic gating variables 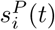 for *P* ∈ {Ext,I} (external and inhibitory currents) are exponential traces of the spike trains of all presynaptic neurons *j* with firing times *t_j_*:

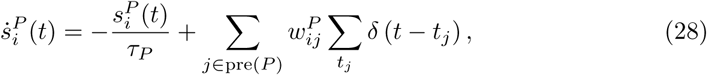

where pre(P) indicates all neurons presynaptic to the neuron *i* for the the connection type *P*. The factors 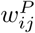 are unit-less synaptic efficacies for the connection from neuron *j* to neuron *i*. For the excitatory gating variables of inhibitory neurons 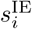 (IE denotes connections from E to *I* neurons) we also use the linear model of Eq. (28) with time constant *τ_IE_* = *τ_E_*.

For excitatory to excitatory conductances, we use a well established model of synaptic short-term plasticity (STP) [48, 91, 92] which provides dynamic scaling of synaptic efficacies depending on presynaptic firing. This yields two additional dynamical variables, the facilitating synaptic efficacy *u_j_*(*t*), as well as the fraction of available synaptic resources *x_j_*(*t*) of the outgoing connections of a presynaptic neuron *j*, which are implemented according to the following differential equation:

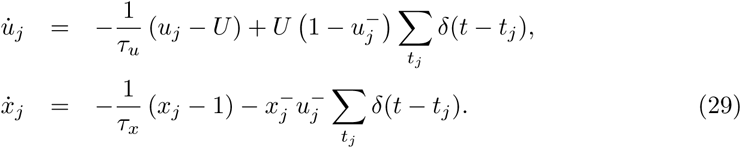

Here, the indices 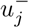 and 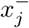 indicate that for the incremental update of the variables upon spike arrival, the values of the respective variables immediately before the spike arrival are used [92]. The variable *U* appears in the equation for *u*(*t*) both as the steady-state value in the absence of spikes and as a scale for the per-spike – intuitively, *U* is thus the first value of *u*(*t*) transmitted on a spike after inactivity of the connection [48].

The dynamics of recurrent excitatory-to-excitatory transmission with STP are then given by gating variables that linearly filter the incoming spikes, which are scaled by facilitation and depression:

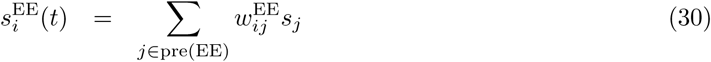

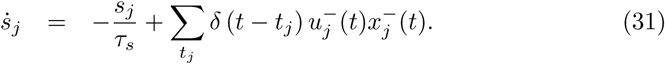

Here, pre(EE) indicates all excitatory neurons that make synaptic connections to the neuron *i*. See the Table in S2 Table for synaptic parameters used in simulations.

This makes the system Eqs. (29) and (31) a spiking variant of the rate-based dynamics of Eq. (3), and 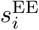 a variable related to the input *J_i_* (cf. Eq. (2)). In *Firing rate approximation* we will make this link explicit.

#### Network connectivity

All connections except for the recurrent excitatory connections are all-to-all and uniform, with unit-less connection strengths set to 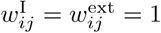 and for inhibitory neurons additionally 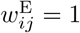. The recurrent excitatory connections are distance-dependent and symmetric. Each neuron of the excitatory population with index *i* ∈ {0,…, *N_E_* − 1} is assigned an angular position 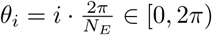. Recurrent excitatory connection weights 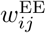 from neuron *j* to neuron *i* are then given by the Gaussian function *w*^EE^(*θ*) as (see the Table in S2 Table for parameters used in simulations):

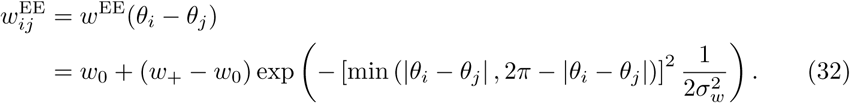

Additionally, for each neuron we keep the integral over all recurrent connection weights normalized, resulting in the normalization condition 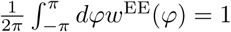. This normalization ensures that varying the maximum weight *w*_+_ will not change the total recurrent excitatory input if all excitatory neurons fire at the same rate. Here, we choose *w*_+_ as a free parameter constraining the baseline connection weight to:

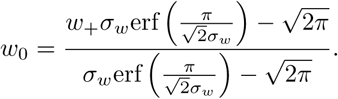

#### Firing rate approximation

We first replace the synaptic activation variables *s^P^*(*V,t*) for *P* ∈ {I, ext} by their expectation values under input with Poisson statistics. We assume that the inhibitory population fires at rates *v_I_*. For the linear synapses this yields

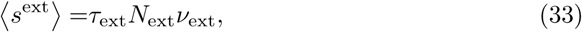

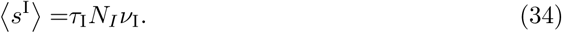

For the recurrent excitatory-to-excitatory synapses with short-term plasticity, we set the differential equations (29) to zero, and also average them over the Poisson statistics. Akin to the “mean-field” model of [48], we average the steady-state values of facilitation and depression separately over the Poisson statistics. This implicitly assumes that facilitation and depression are statistically independent, with respect to the distributions of spike times – while this is not strictly true, the approximations work well, as has been previously reported [48]. This allows a fairly straightforward evaluation of the mean steady-state value of the combined facilitation and depression variables 〈*u_j_x_j_*〉, under the assumption that the neuron *j* fires at a mean rate *v_j_* with Poisson statistics, and yields rate approximations of the steady-state values similar to Eq. (12):

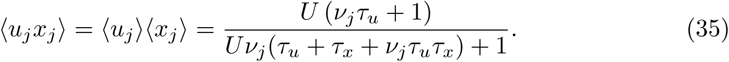

We now assume that the excitatory population of *N_E_* neurons fires at the steady-state rates *ϕ_j_* (0 ≤ *j* < *N*). To calculate the synaptic activation of excitatory-to-excitatory connections 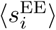, we set Eq. (30) to zero, and average over Poisson statistics (again neglecting correlations), which yields 〈*s_j_*〉 = *τ_E_*〈*u_j_x_j_*〉*ϕ_j_* and 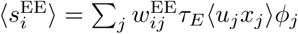. Let the the normalized steady-state input *J_i_* be:

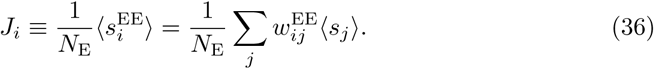

The steady-state input Eq. (36) links the general framework of Eq. (2) to the spiking network. The additional factor 1/*N*_E_ is introduced to make the scaling of the excitatory-to-excitatory conductance with the size of the excitatory population *N*_E_ explicit, which will be used in *System size scaling*. To see this, we assume that the excitatory conductance scale of excitatory neurons gEE is scaled such that the total conductance is invariant under changes of *N_E_* [93]: 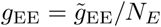, for some fixed value 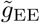. This yields the total excitatory-to-excitatory conductance 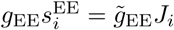 with *J_i_* as introduced above, where the scaling with *N_E_* is now shifted to the input variable *J_i_*.

For the synaptic activation of excitatory to inhibitory connections, we get the mean activations:

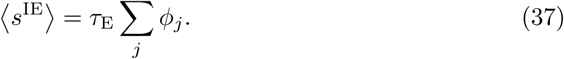

We then follow [49] to reduce the differential equations of Eq. (27) to a dimensionless form. The main difference consists in the absence of the voltage dependent NMDA conductance, which is achieved by setting the two associated parameters *β* → 0, *γ* → 0 in [49], to arrive at:

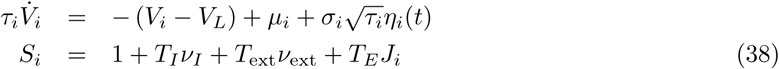

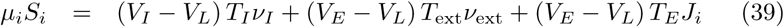

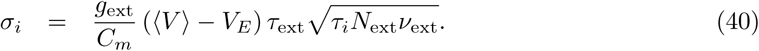

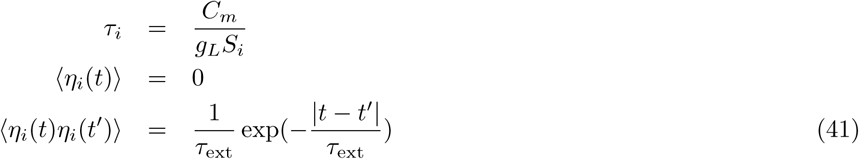

where 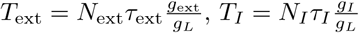 are effective timescales of external and inhibitory inputs, and 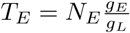 is a dimensionless scale for the excitatory conductance. Here, *μ_i_* is the bias of the membrane potential due to synaptic inputs, and *σ_i_* measures the scale of fluctuations in the membrane potential due to random spike arrival approximated by the Gaussian process *η_i_*.

The mean firing rates *F* and mean voltages 〈*V_i_*〉 of populations of neurons governed by this type of differential equation can then be approximated by:

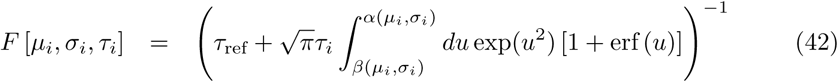

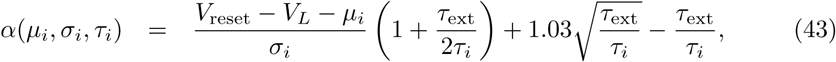

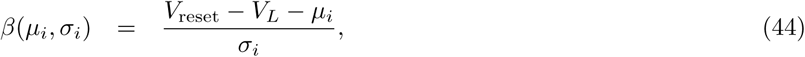

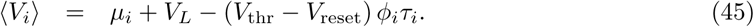

#### Derivatives of the rate prediction

Here we calculate derivatives of the input-output relation (Eq. (42)) that will be used below in *Frozen noise*.

The expressions for drift and diffusion (see *Analysis of drift & diffusion with STP*) contain the derivative 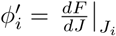 of the input-output relation *F* (Eq. (42)) with respect to the recurrent excitatory input; *J_i_*. Note, that *F* depends on *J_i_* through all three arguments *μ_i_,σ_i_* and *τ_i_*. First, we define *X*(*u*) ≡ exp(*u*^2^) [1 + erf(*u*)], and the shorthand *F_i_* = *F* [*μ_i_,σ_i_,τ_i_*]. The derivative can then be readily evaluated as (to shorten the notation in the following, we skip noting the evaluation points for derivatives in the following):

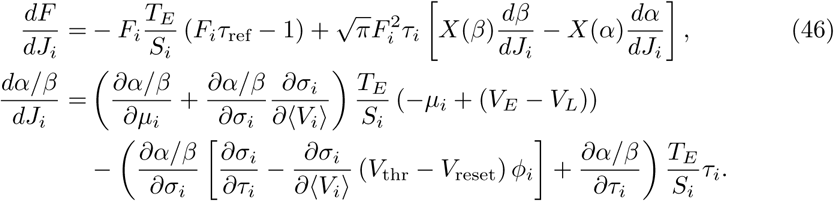

where *α*/*β* stands as a placeholder for either function, and the expressions for *α* and *β* are given in Eqs. (43)-(44).

A second expression involving the derivative of Eq. (42) is 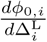 which appears in the theory when estimating firing rate perturbations caused by frozen heterogeneities in the leak potentials of excitatory neurons (see Eq. (54)). The resulting derivatives are almost similar, which can be seen by the fact that replacing 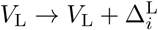 in Eq. (27) only leads to an additional term 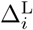 in Eq. (39). Thus, for neuron *i* the derivative can be evaluated to

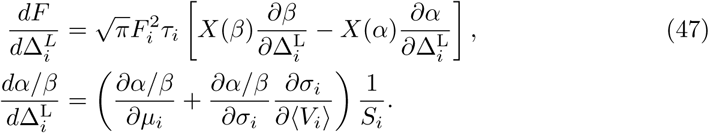

In practice, given a vector *ϕ*_*i*,0_ of firing rates in the attractor state, as well as the mean firing rate of inhibitory neurons *v_I_*, we evaluate the right hand side of Eq. (46) and Eq. (47) by replacing *F_i_* → *ϕ*_*i*,0_. This allows efficiently calculating the derivatives without having to perform any numerical integration. The two terms will be exactly equal if *ϕ*_0,*i*_ is a self-consistent solution of Eq. (42) for firing rates of the excitatory neurons across the network. We used numerical estimates of *ϕ*_0,*i*_ and *v_I_* that were measured from simulations and were very close to firing-rate predictions for all networks we investigated.

#### Optimization of network parameters

We used an optimization procedure [94] to retune network parameters to produce approximately similar bump shapes as the parameters of short-term plasticity are varied. Briefly, we replace the network activity *ϕ_j_* in the total input *J_i_* of Eq. (36) by a parametrization

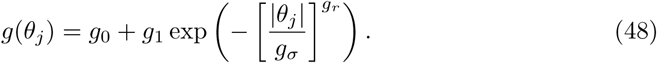

Approximating sums 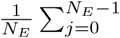 with integrals 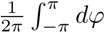 we arrive at

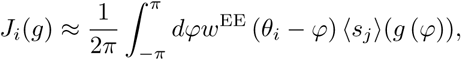

where *J_i_*(*g*) indicates that the total input depends on the parameters *g*_0_, *g*_1_, *g_σ_, g_r_* of the parametrization *g*.

We then substitute this relation in Eq. (42) to arrive at a self-consistency relation between the parametrized network activity *g*(*θ_i_*) at the position of neuron *i* and the firing-rate *F* predicted by the theory:

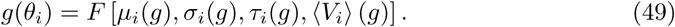

As for the input *J_i_*(*g*), we indicate the dependence of quantities on the parameters of the parametrization *g* by appending (*g*). The explicit dependence of the voltage 〈*V_i_*〉(*g*) on *g* is obtained by additionally substituting *ϕ_i_* → *g*(*θ_i_*) in Eq. (45).

We then optimized networks to fulfill Eq. (49). First, we imposed the following targets for the parameters of *g*: *g*_0_ = 0.1Hz, *g*_1_ = 40Hz, *v*_*E*,basal_ = 0.5Hz, *v*_*I*,basal_ = 3Hz. For all networks we chose *w*_+_ = 4.0, *g_r_* = 2.5. The following parameters were then optimized: *v_I_, g_σ_, g*_EE_ (excitatory conductance *g_E_* on excitatory neurons); *g*_IE_ (excitatory conductance *g*_E_ on inhibitory neurons); *g*_EI_ (inhibitory conductance *g_I_* on excitatory neurons); *g*_II_ (inhibitory conductance *g*_II_ on inhibitory neurons). The basal firing rates (firing rates in the uniform state of the network, prior to being cued) yielded two equations from Eq. (49) by setting *w*_+_ = 1. This left 4 free parameters, which were constrained by evaluating Eq. (49) at 4 points as described in [94]. The basal firing rates were chosen to be fairly low to make the uniform state more stable (as in [44]). This procedure does not yield a fixed value for *g_σ_*, since *g_σ_* is optimized for and is not set as a target value. We thus iterated the following until a solution was found with *g_σ_* ≈ 0.5: a) change the width of the recurrent weights *w_σ_*; b) optimize network parameters as described here; c) optimize the expected bump shape for the new network parameters to predict *g_σ_*. The resulting parameter values are given in Table in S2 Table.

### Frozen noise

#### Random and heterogeneous connectivity

Introducing random connectivity, we replace the recurrent weights in Eq. (36) by:

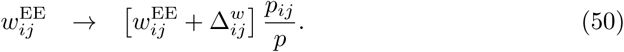

Here, *p_ij_* ∈ {0,1} are Bernoulli variables, with *P*(*p_ij_* = 1) = *p*, where the connectivity parameter *p* ∈ (0,1] controls the overall sparsity of recurrent excitatory connections. For *p* = 1 the entire network is all-to-all connected. Additionally, we provide derivations for additive synaptic heterogeneities 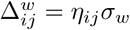 (as in [38]), where {*η_ij_*|1 ≤ *i, j* ≤ *N_E_*} are independent, normally distributed random variables with zero mean and unit variance. We did not investigate this type of heterogeneity in the main text, since increasing lead *σ_w_* to a loss of the attractor state before creating large enough directed drifts to be comparable to the other sources of frozen noise considered here – most of the small effects were “hidden” behind diffusive displacement [81]. Nevertheless, we included this case in the analysis here for completeness.

Let the center position of the bump be 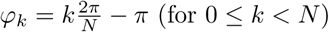. Subject to the perturbed weights, the recurrent steady-state excitatory input *J_i_*(*φ_k_*) Eq. (36) to any excitatory neuron can be written as the unperturbed input *J*_0,*i*_(*φ_k_*) plus an additional input 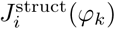 arising from the perturbed connectivity. Note that the synaptic steady-state activations *s*_0,*j*_(*φ_k_*) change with varying bump centers – in the following, we denote 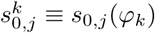:

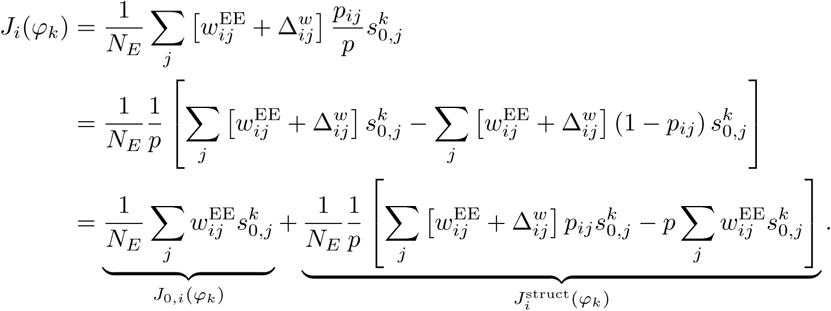

Note that *J*_0,*i*_(*φ_k_*) is an index-shifted version of the steady-state input: *J*_0,*i*_(*φ_k_*) = *J*_0,*i*−*k*_. However, such a relation does not hold for 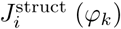, since the random numbers *p_ij_* will change the resulting value for varying center positions.

We calculate the firing rate perturbations *δϕ_i_*(*φ_k_*) resulting from the additional input by a linear expansion around the steady-state firing rates *ϕ*_0,*i*_(*φ_k_*) → *ϕ*_0,*i*_(*φ_k_*) + *δϕ_i_*(*φ_k_*). These evaluate to:

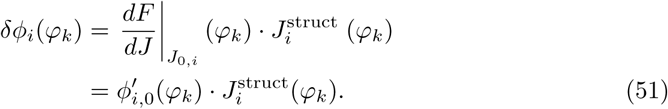

See *Derivatives of the rate prediction* for the derivation of the function 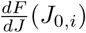 for the spiking network used in the main text.

In the sum of Eq. (7), we keep the firing rate profile 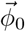 centered at *φ*_0_ while calculating the drift for varying center positions. To accommodate the shifted indices resulting from moving center positions, we re-index the summands to yields the perturbations *ϕ*_0,*i*_ → *ϕ*_0,*i*_ + Δ*ϕ_i_*(*φ_k_*) used there:

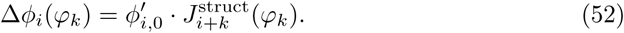

#### Heterogeneous leak reversal potentials

We further investigated random distributions of the leak reversal potential *V*_L_. These are implemented by the substitution

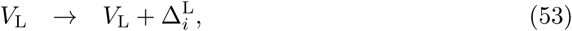

where the 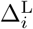 are independent normally distributed variables with zero mean, i.e. 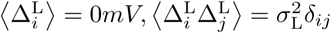. The parameter *σ*_L_ controls the standard deviation of these random variables, and thus the noise level of the leak heterogeneities.

Let 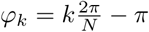 for 0 ≤ *k* < *N* be the center position of the bump. First, note that the heterogeneities 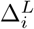 do not depend on the center position *φ_k_*, since they are single neuron properties. As in the last section, we calculate the firing rate perturbations *δϕ_i_*(*φ_k_*) resulting from the additional input by a linear expansion around the steady-state firing rates *ϕ*_0,*i*_(*φ_k_*) → *ϕ*_0,*i*_(*φ_k_*) + *δϕ_i_*(*φ_k_*):

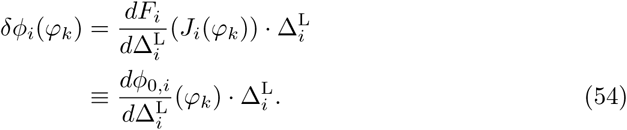

Here, 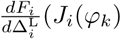 is the derivative of the input-output relation of neuron *i* in a bump centered at *φ_k_*, with respect to the leak perturbation. We introduced 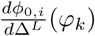 as a shorthand notation for this derivative, since it is evaluated at the steady-state input *J*_*i*,0_(*φ_k_*). For the spiking network of the main text, this is derived in *Derivatives of the rate prediction*.

In the sum of Eq. (7), we keep the firing rate profile 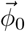 centered at *φ*_0_ while calculating the drift for varying center positions. As in the last section, we re-index the sum to yield the perturbations *ϕ*_0,*i*_ → *ϕ*_0,*i*_ + Δ*ϕ_i_*(*φ_k_*) used there:

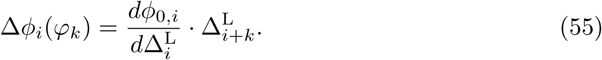

#### Squared field magnitude

Using the equation of the drift field in Eq. (7), and the firing rate perturbations Eqs. (51)-(54), it is straight forward to see that for any center position *φ* the expected drift field averaged over the noise parameters is 0, since all single firing rate perturbations vanish in expectation. In the following we calculate the variance of the drift field averaged over noise realizations, which turns out to be additive with respect to the two noise sources.

We begin by calculating the correlations between frozen noises caused by random connectivity and leak heterogeneities. For the Bernoulli distributed variables *p_ij_* it holds that 〈*p_ij_*〉 = *p*, 〈*p_ij_p_lk_*〉 = *δ_il_δ_jk_p* + (1 − *δ_il_δ_jk_*)*p*^2^. For the other independent random variables it holds that 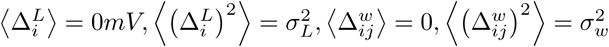. Again, the weight heterogeneities 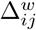, are only included for completeness – all analyses of the main text assume that *σ_w_* =0.

For the correlations between the perturbations we then know that (for brevity, we omit the dependence on the center position *φ*):

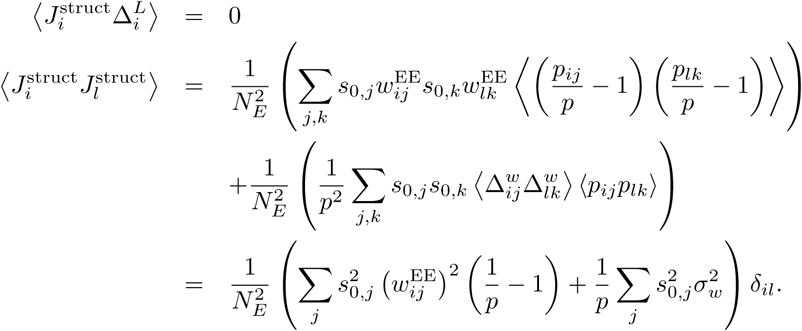

Starting from Eq. (7), we use as a firing rate perturbation the sum of firing rate perturbations from both Eq. (51) and Eq. (54). With the pre-factor 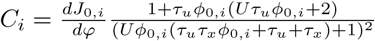, the expected squared field averaged over ensemble of frozen noises is then:

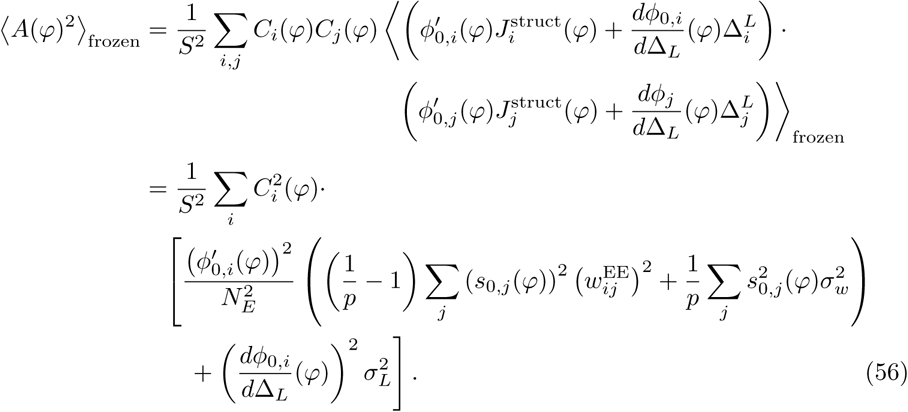

One can see directly that the two last terms are invariant under shifts of the bump center *φ*, since these introduce symmetric shifts of the indexes *i*. Similarly, it is easy to see that the first term is also invariant. Let *φ*′ be shifted to the right by one index from *φ*. It then holds that:

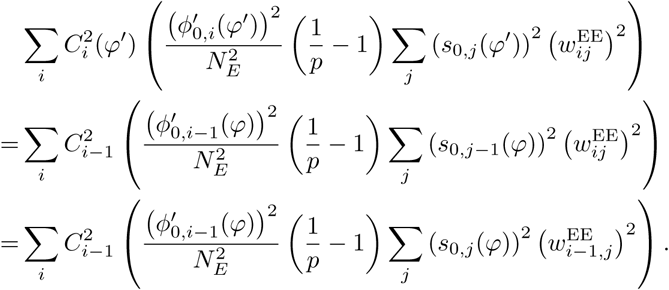

The final equation holds since, in ring-attractor networks, 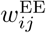 consists of index-shifted rows of the same vector (see e.g. *Network connectivity* for the spiking network weights).

In summary, 〈*A*(*φ*)^2^〉_frozen_, will evaluate to the same quantity 〈*A*^2^〉_frozen_. for all center positions *φ*. In the main text, we use this fact to estimate 〈*A*^2^〉_frozen_ from simulations, by additionally averaging over the all center positions and interchanging the ensemble and positional averages:

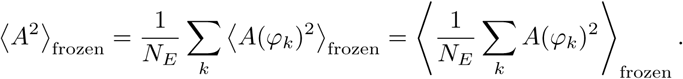

Thus, we can compare the value of 〈*A*^2^〉_frozen_ to the mean squared drift field over all center positions, averaged over instantiations of noises.

#### System size scaling

Generally, sums over the discretized intervals [−*π, π*) as they appear in Eqs. (5) and (7) will scale with the number *N* chosen for the discretization of the positions on the continuous ring 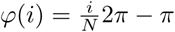. Consider two discretizations of the ring, partitioned into *N*_1_ and *N*_2_ uniformly spaced bins of width 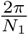 and 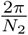. We can then approximate integrals over any continuous (Riemann integrable) function *f* on the ring by the two Riemann sums:

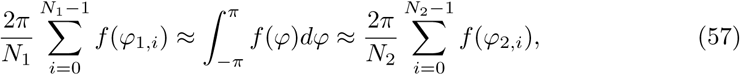

where, 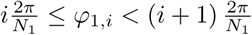 (for *N*_2_ and *φ*_2,*i*_ analogously) are points in the bins [95].

Numerical quantities for the results of the main text have been calculated for *N_E_* = 800. In the following we denote all of these quantities with an asterisk (*). To generalize these results to arbitrary system size *N*, we replace sums over *N* bins bye scaled sums over *N_E_* bins using the relation Eq. (57):

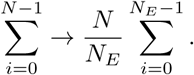

First, we find that the normalization constant scales as 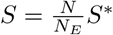, and thus (dots indicate the summands, which are omitted for clarity) for the diffusion strength B (cf. Eq. (5)):

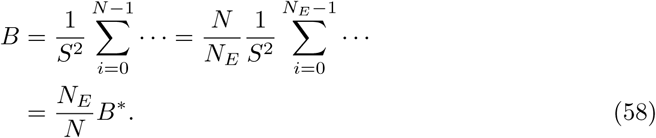

For the drift magnitude we turn to the expected squared drift magnitude calculated earlier (cf. Eq. (56)), for which we find that (setting *σ_w_* → 0 for simplicity, as throughout the main text):

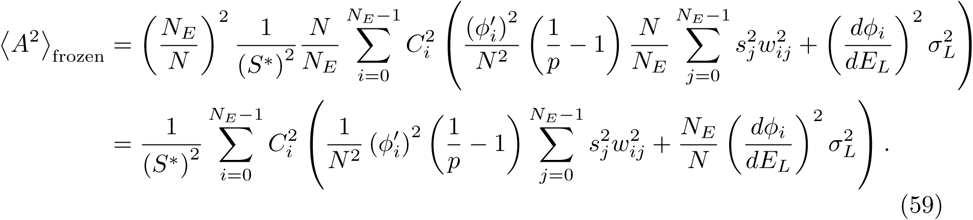

Note, that we could not resolve this scaling in dependence of 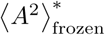, since the two sources of frozen noise (connectivity and leak heterogeneity) show different scaling with *N*.

### Numerical methods

#### Spiking simulations

All network simulations and models were implemented in the NEST simulator [96]. Neuronal dynamics are integrated by the Runge-Kutta-Fehlberg method as implemented in the GSL library [97] (gsl_odeiv_step_rkf45) – this forward integration scheme is used in the NEST simulator for all conductance-based models (at the time of writing). The short-term plasticity model is integrated exactly, based on inter-spike intervals. Code for network simulations will be made available upon publication at https://github.com/EPFL-LCN/pub-seeholzer2018.

##### Simulation protocol

In all experiments (except those involving bi-stability, see below) spiking networks were simulated for a transient initial period of *t*_initial_ = 500*ms*. To center the network in an attractor state at a given angle −*π* ≤ *φ* < *π*, we gave an initial cue signal by stimulating 0.1 · *N_E_* neurons centered at *φ* by strong excitatory input mediated by additional Poisson firing onto AMPA receptors (1*s*, 2kHz) with connections scaled down by a factor of *g*_signal_ = 0.5. The external input ceased at *t* = *t*_off_ = 1.5*s*. For simulations to estimate the diffusion we simulated until *t*_max_ = 15*s*, yielding 13.5*s* of delay activity after the cue offset. For simulations to estimate drift we set *t*_max_ = 8*s*, yielding 6.5*s* of delay activity after the cue offset.

For simulations exploring the bi-stability between the uniform state and a bump state (Fig. 1B1), we added an additional input prior to the spontaneous state. We stimulated simultaneously 20 excitatory neurons around 4 equally spaced cue points each (80 neurons in total, 500*ms*, 1.5kHz, AMPA connections scaled by a factor *g*_signal_ = 2). This was applied to settle networks into the uniform state more stably – without this perturbation, networks sometimes approached the bump state after being uniformly initialized. In both figures, we show population activity only after this initial stimulus was applied.

##### Estimation of centers and mean bump shapes

To estimate centers of bump states, simulations were run until *t* = *t*_max_ and spikes were recorded from the excitatory population and converted to firing rates by convolving them with an exponential kernel (*τ* = 100*ms*) [98] and then sampled at resolution 1ms. This results in vectors of firing rates *v_j_*(*t*), 0 ≤ *j* ≤ *N_E_* − 1 for every time *t*. We calculated the population center *φ*(*t*) for time *t* by measuring the phase of the first spatial Fourier coefficient of the firing rates. This is given by 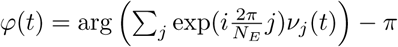. For all analyses below, we identify *t* = 0 to be the time *t* = *t*_off_ of the initial cue.

To measure the mean bump shapes, we first rectified the vectors *v_j_*(*t*) for every *t* by rotating the vector until *φ*(*t*) = 0. We then sampled the rectified firing rates starting from 1*s* after cue offset at intervals of 20ms, which were used to calculate the mean firing rates. S1 Fig shows mean rates for each simulation averaged over the ~ 1000 repetitions performed in the diffusion estimation (below).

##### Exclusion of bump trajectories

Sometimes bump trajectories would leave the attractor state and return to the uniform state. We identified these trajectories in all experiments by identifying maximal firing rates across the population that dropped below 10Hz during the delay period. The such identified repetitions were excluded from the analyses, which occurred mostly in networks with no facilitation for *τ_x_* = 150*ms, τ_u_* = 650*ms*: at *U* = 1, we excluded 222/1000 repetitions from the diffusion estimation, while for all other *U* ≤ 0.8 at most 15/1000 were excluded. Increasing the depression time constant also lead to less stable attractor states: for *τ_x_* = 200*ms, τ_u_* = 650*ms* and *U* = 0.8, we had to exclude 250/1000 repetitions. During the simulations for drift estimation, we observed that frozen noise also leads to less stable bumps under weak facilitation for random and sparse connectivity (*p* ≪ 1) and high leak variability (*σ_L_* ≫ 0).

##### Diffusion estimation

Diffusion was estimated for each combination of network parameters by simulating 1000 repetitions (10 initial cue positions, 100 repetitions each) of 13.5*s* of delay activity. Center positions *φ_k_*(*t*) were estimated for each repetition *k* as described above. We then calculated for each repetition the offset relative to the position at 500*ms* by Δ*φ_k_*(*t*) = *φ_k_*(*t* − 500*ms*) − *φ_k_*(500*ms*), effectively discarding the first 500*ms* after cue-offset. The time-dependent variance of *K* repetitions (excluding those repetitions in which the bump state was lost, see above) was then calculated as 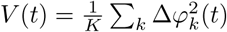. The diffusion strength can then be estimated from the slope of a linear least-squares regression (using the Scipy method *scipy.stats.linregress* [99]) to the variance as a function of time: *V*(*t*) ≈ *D*_0_ + *D* · *t*, where the intercept *D*_0_ is included to account for initial transients. We estimated confidence intervals by bootstrapping [100]: sampling K elements out of the K repetitions with replacement (5000 samples) and estimating the confidence level of 0.95 by the bias corrected and accelerated bootstrap implemented in *scikits-bootstrap* [101]. As a control, we calculated confidence intervals for *D* additionally by Jackknifing: after building a distribution of estimates of D on *K* one-left-out samples of all repetitions, the standard error of the mean can be calculated and is multiplied by 1.96 to obtain the 95% confidence interval [102] – confidence intervals obtained by this method were almost indistinguishable from confidence intervals obtained by bootstrapping.

##### Drift estimation

Drift was estimated numerically for each combination of network and frozen noise parameters by simulating 400 repetitions (20 initial cue positions, 20 repetitions each) of 6.5*s* of delay activity. Centers positions *φ_k_*(*t*) were estimated for all *K* repetitions (excluding those repetitions in which the bump state was lost, see above) as explained above. We then computed displacements in time by computing a set of discrete differences

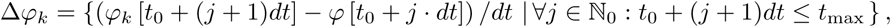

where we chose *dt* = 1.5*s* and *t*_0_ ∈ {500*ms*, 700*ms*, 900*ms*,…, 1900*ms*}. All differences are calculated with periodic boundary conditions on the circle [−*π, π*), i.e. the maximal difference was *π*/*dt*. We then calculated a binned mean (100 bins on the ring, unless mentioned otherwise) of differences calculated for all *K* trajectories, to approximate the drift-fields as a function of positions on the ring.

#### Mutual information measure

We are estimating the mutual information between a set of initial positions *x* ∈ [0, 2*π*) and associated final positions *y*(*x*) ∈ [0, 2*π*) of the trajectories of a continuous attractor network over a fixed delay period of *T*. For our results, we take *T* = 6.5*s*. We constructed binned and normalized histograms (with bin size *n* = 100, but see below) as approximate probability distributions of initial positions 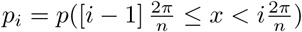 and all final positions 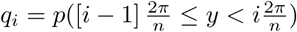 (with bins indexed by 1 ≤ *i* ≤ *n*), as well as the bivariate probability distribution 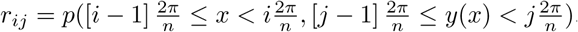.

Using these, we can calculate the mutual information as [54, 55] 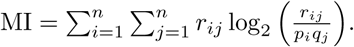. Note, that the sum effectively counts only nonzero entries of *r_ij_* (trajectories that started in bin *i* and ended in bin *j*): these imply that *p_i_* ≠ 0 (a trajectory started in bin *i*) and *q_j_* ≠ 0 (a trajectory ended in bin *j*), which makes the sum well defined. Although the value of MI depends on the number of bins *n*, in Fig. 5 and Fig. 6 we normalize MI to that of the reference network (*U* = 1, no frozen noise, see *Short-term plasticity controls memory retention*), which leaves the resulting plot nearly invariant under a change of bin numbers.

#### Numerical integration of Langevin equations

Numerically integration of the homogeneous Langevin equations (Eq. (4)) describing drift and diffusion of bump positions *φ* ∈ [−*π, π*) (with circular boundary conditions) has been implemented as a *C* extension in Cython [103] to the Python language [104]. Since the drift fields *A*(*φ*) are estimated on a discretization of the interval [−*π, π*) into *N* bins, we first interpolate drift fields *A* given as *N* discretized values to obtain continuous fields – interpolations are obtained using cubic splines on periodic boundary conditions using the class *gsl_interp-cspline_periodic* of the Gnu Scientific Library [97].

For forward integration of the Langevin equation Eq. (4) from time *t* = 0, we start from an initial position *φ*_0_ = *φ*(*t* = 0). Given a time resolution *dt* (unless otherwise stated we use *dt* = 0.1*s*) and a maximal time *t*_max_ we repeat the following operations until we reach *t* = *t*_max_:

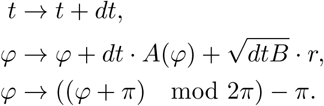

Here, for each iteration *r* is a random number drawn from a normal distribution with zero mean and unit variance (〈*r*〉 =0 and 〈*r*^2^〉 = 1). The last step is performed to implement the circular boundary conditions on [−*π, π*).

Code implementing this numerical integration scheme will be made available upon publication at https://github.com/EPFL-LCN/pub-seeholzer2018-langevin.

#### Distractor analysis

For the distractor analysis in Fig. 7, we let 40 neurons centered at the distractor position 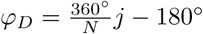 fire at rates increased by 20Hz, yielding a vector of firing rate perturbations Δ*ϕ*_0,*i*_ = 20Hz if |*i* − *j*| ≤ 20 and Δ*ϕ*_0,*i*_ = 0Hz otherwise. The vectors Δ*ϕ*_0,*i*_ foreach distractor position *φ_D_* are then used in Eq. (7) to calculate the corresponding drift fields. To calculate the final position *φ*_1_ after 250ms of presenting the distractor, we generate 1000 trajectories starting from *φ*_0_ = 0 by integrating the Langevin equation Eq. (4) for 250ms (*dt* = 0.01), the final positions of which are used to measure mean and standard deviation of *φ*_1_. For the broader bump in Fig. 7D, we stretched (and interpolated) the firing rates *ϕ*_0_ as well as the associated vectors *J*_0_ and 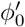 along the x-axis to obtain vectors for bumps of the desired width, and then re-calculated the values of 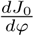.

## Acknowledgments

The authors thank Tilo Schwalger and Johanni Brea for helpful discussions and feedback. Research was supported by the European Union Seventh Framework Program (FP7) under grant agreement no. 604102 (Human Brain Project, M.D.) and by the Swiss National Science Foundation (200020_147200, A.S.).

## Supporting information

**S1 Fig. Spiking networks produce similar stable firing rate profiles across parameters.** For each choice of short-term plasticity parameters *U, τ_u_*, and *τ_x_*, we tuned the recurrent conductances (*g*_EE_, *g*_EI_, *g*_IE_, *g*_II_) and the width *σ_w_* of the distance-dependent weights (cf. Eq. (32)) such that the “bump” shape of the stable firing rate profile is close to a generalized Gaussian 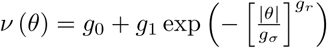 with parameters *g*_0_ = 0.1Hz, *g*_1_ = 40.0Hz, *g_σ_* = 0.5, *g_r_* = 2.5. See *Optimization of network parameters* in Materials and Methods for details, S2 Table for parameter values after tuning, and S1 Table for parameters that stay constant. **A** After tuning, the resulting firing rate profiles for different parameter values of *U* and *τ_u_* are very similar. Averaged mean firing rates in bump state, measured from ~ 1000 spiking simulations. **A1-A3** Remaining slight parameter-dependent changes of bump shapes, measured by fitting the generalized Gaussian *v*(*θ*) to the measured firing rate profiles displayed in A. **A1** Top firing rate *g*_1_. **A2** Half-width parameter *g_σ_*. **A3** Sharpness parameter *g_r_*. **B** and **B1-B3** Same as in A and A1-A3, for additional variation of the depression time scale *τ_x_*.

**S2 Fig. Theoretical prediction of diffusion strength as a function of STP parameters.** All color values display diffusion magnitude estimated from *B* in Eq. (4) with bump shape estimated from the reference network (*U* = 1, *τ_x_* = 150ms, compare 3B,C, dashed lines). Units of color values are 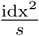 with values of level lines as indicated. **A** Diffusion as function of facilitation *U* and depression time constant *τ_x_*. Facilitation time constant was *τ_u_* = 650*ms*. **B** Diffusion as function of facilitation *U* and facilitation time constant *τ_u_*. Depression time constant was *τ_x_* = 150*ms*. **C** Diffusion as function of depression time constant *τ_x_* and facilitation time constant *τ_u_*. Facilitation *U* was *U* = 0.5.

**S3 Fig. Comparison of theoretically predicted fields to simulations. A** Averaged root mean square error (RMSE) between predicted fields (Eq. (7)) and fields extracted from simulations (mean over 18-20 networks, error bars show 95% confidence of the mean). Both frozen noise parameters (*σ*_L_ and 1 − *p*) are plotted on the same x-axis. **B** Normalized RMSE: each RMSE is normalized by the range (max − min) of the joint data of simulated and predicted fields it is calculated on. Colors as in A. **C** Average RMSE (same data as in A) plotted as a function of the mean expected field magnitude (estimated separately for each network, then averaged). Colors as in A. **D** Worst (top) and best (bottom) match between predicted field (blue line) and field extracted from simulations (black line) of the group with the largest mean RMSE in panels A, C (*U* = 1, 1 − *p* = 0.75). Shaded areas show 1 standard deviation of points included in the binned mean estimate (100 bins) of the extracted field.

**S4 Fig. Mutual information normalized to compare slopes.** Same data as in Fig. 5B, but MI is normalized to the average MI of each spiking network without heterogeneities (leftmost dot for each green, orange, and blue group of curves/dots), making explicitly visible the change in slope of the drop-off as heterogeneity parameters are increased. Dashed lines connect the means, for visual guidance.

**S5 Fig. Theoretical predictions of working memory stability.** All panels show theoretically predicted expected displacement over 1 second (Eq. (11)) for networks with random and sparse connections (*p* = 0.12) and leak reversal potential heterogeneity (*σ_L_* = 1.7*mV*). White lines show displacement contour lines for 1, 2 and 5 deg. **A** Displacement as a function of the facilitation time constant *τ_u_* and facilitation *U* for *τ_x_* = 150*ms* and *N* = 5000.**B** Displacement as a function of system size and facilitation *U* for *τ_x_* = 150*ms* and *τ_u_* = 650*ms*. **C-D** Displacement as a function of depression time constant *τ_x_* and facilitation *U* for *N* = 5000 (C) and *N* = 20000 (D). In both panels *τ_u_* = 650*ms*.

**S6 Fig. Dependence of diffusion strength B on shape parameters.** Diffusion was calculated from Eq. (5) with bump solutions 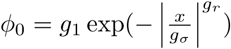. The values of 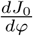 and 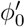 were calculated by fitting and extrapolating (linearly, for *ϕ*_0_ > 40.31Hz) curves 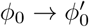 and *ϕ*_0_ → *J*_0_ that were obtained from the numerical values extracted for *g*_1_ = 40.31Hz, *g_σ_* = 0.51 by theory (see *Firing rate approximation* in Materials and Methods). Thus, any nonlinearity or saturation of the inputs and input-output relation for *ϕ*_0_ > 40.31Hz was not included. This approximate analysis shows that the major dependence of the diffusion expected in the system is on the bump width *g_σ_*, although a minor dependence on *g*_1_ is seen.

**S7 Fig. Short-term plasticity does not affect spiking statistics.** Mean firing rate, coefficient of variation of the inter-spike interval distribution (CV), and local CV (*CV*_2_ [89]) for two attractor networks with different STP parameters. All measures were computed on spike-trains measured over a period of 4*s*, recorded 500*ms* after offset of the external input which was centered at angle 0. Across STP parameters, networks display similarly reduced CVs for increased mean firing rates, leading to large CVs for neurons located in the flanks of the firing rate profile and low CVs for neurons located near the center. **A** Networks with large diffusion coefficient (*U* = 0.8, *τ_u_* = 650*ms, τ_x_* = 200*ms*) that underwent non-stationary diffusion during the recording of spikes: the measured mean firing rates (gray line) differ visibly from the firing rates estimated after centering the firing rate distribution at each point in time. Due to this non-stationarity, CVs at intermediate firing rates appear elevated, while the local CV (*CV*_2_) shows values close to stationary networks (see B). **B** The same network as in A, with strong facilitation (*U* = 0.1). Reduced diffusion leads to a nearly stationary firing rate profile, and coincident CV and *CV*_2_ measures.

### S1 Supporting Information. Detailed mathematical derivations

**S1 Table. Parameters for spiking simulations.** Parameter values are modified from [11] and [49]. For recurrent conductances see the table in S2 Table.

**S2 Table. Conductance and connectivity parameters for spiking simulations.** For all networks we set *w*_+_ = 4.0. Recurrent conductance parameters are given for combinations of short-term plasticity parameters according to the following notation. *g*_EE_: excitatory conductance *g*_E_ on excitatory neurons; *g*_IE_: excitatory conductance *g*_E_ on inhibitory neurons; *g*_EI_: inhibitory conductance *g*_I_ on excitatory neurons; *g*_II_: inhibitory conductance *g*_I_ on inhibitory neurons.

## Footnotes

1 In Brownian motion, the *diffusion constant* is usually defined as *D* = *B*/2.

